# Bacterial cell-free DNA profiling reveals co-elevation of multiple bacteria in newborn foals with suspected sepsis

**DOI:** 10.1101/2024.10.31.620104

**Authors:** Li-Ting Chen, Emmy Wesdorp, Myrthe Jager, Esther W. Siegers, Mathijs J.P. Theelen, Nicolle Besselink, Carlo Vermeulen, Aldert L. Zomer, Els M. Broens, Jaap A. Wagenaar, Jeroen de Ridder

**Affiliations:** Center for Molecular Medicine, University Medical Center Utrecht, Utrecht University, 3584 CX Utrecht, The Netherlands; Oncode Institute, 3521 AL, Utrecht, the Netherlands; Department of Clinical Sciences, Faculty of Veterinary Medicine, Utrecht University, 3584 CM Utrecht, The Netherlands; Department of Biomolecular Health Sciences, Faculty of Veterinary Medicine, Utrecht University, 3584 CL Utrecht, the Netherlands; Wageningen Bioveterinary Research, 8221 RA Lelystad, The Netherlands

## Abstract

**Background:** Sepsis is the leading cause of death in newborn foals. This study investigates whether cell-free DNA (cfDNA) sequencing can enhance bacterial pathogen detection in foals with suspected sepsis and addresses existing knowledge gaps and diagnostic challenges.

**Methods:** We developed a **f**oal <u>cf</u>DNA sequencing for **b**acteria **i**dentification (<u>cf</u>**FBI**) workflow, integrating wetlab and computational protocols to detect increased bacterial cfDNA abundance in blood. Specifically, cfFBI focusses on enriching bacterial cfDNA molecules and preventing false positive bacterial identifications. cfFBI was applied to blood samples of 25 hospitalized foals categorized according to the neonatal Systemic Inflammatory Response Syndrome (nSIRS) criteria and 7 healthy foals.

**Results:** cfDNA levels of potential sepsis-causing bacterial genera were elevated in all 11 nSIRS-positive foals compared to healthy foals (n=7), and in 8/11 (72.7%) when compared to both nSIRS-negative (n=4; nSIRS=0) and healthy foals, with multiple genera elevated in 5/11 (45.5%). The total cfDNA concentration, bacterial cfDNA fraction and bacterial diversity were not different between the foal groups. However, nSIRS-positive foals showed significantly different end-motifs in host chromosomal cfDNA, and a decrease in host mitochondrial cfDNA fraction.

**Conclusions:** This study is the first to demonstrate that cfDNA sequencing in blood samples from newborn foals enables detection of pathogenic bacteria and can help identify novel host-related sepsis biomarkers. The elevated presence of multiple sepsis-causing genera in nSIRS-positive foals and the difference in end-motif, suggests that multibacterial elevation may be more common than previously thought. These findings indicate that cfDNA sequencing holds promise as a future diagnostic tool for identifying sepsis in newborn foals.

## Introduction

Sepsis is defined as “a life-threatening organ dysfunction caused by a dysregulated host response to infection”, hallmarked by the systemic inflammatory response syndrome (SIRS) and often caused by a bacterial infection ^1,2^. SIRS arises when pathogen- and damage-associated molecular patterns (PAMPs and DAMPs), as well as neutrophil extracellular traps (NETs), are recognized by the immune system ^3,4^. Dysregulated innate and adaptive immune responses, in combination with overactivation of the coagulation system can result in multiple organ dysfunction, followed by multiple organ failure, and ultimately resulting in death ^5–7^. In newborn foals, sepsis stemming from a bacterial infection is an important cause of morbidity and mortality during the first week of life ^8–10^. Due to the rapid progression of sepsis, early recognition, prompt identification of the causative bacterial pathogen, and timely initiation of effective antimicrobial therapy are critical for improving survival rates^11^.

Despite being a common cause of death in newborn foals, knowledge gaps persist regarding the pathogenesis, diagnosis, and treatment of sepsis. For instance, multiple bacteria are known to co-occur in human sepsis patients ^12,13^, and a similar phenomenon is likely in newborn foals ^11,14^. However, traditional culture methods often lack the sensitivity to detect such co-infections. Additionally, when multiple bacteria are cultured from a single sample it is frequently dismissed as contamination in clinical settings. A further complication in understanding sepsis pathogenesis and diagnostics is that newborn foals absorb immunoglobulins from the colostrum over the gastrointestinal barrier, during which bacteria (including those that can cause sepsis) can also enter the bloodstream ^15^. Although transient bacteremia is a normal physiological process, it remains unclear why in some foals this might lead to SIRS and sepsis, while in the majority it does not.

Early sepsis diagnosis and timely identification of the causative microbe(s) remains challenging in foals with current diagnostic tools. To aid prompt identification of foals at risk for sepsis, several scoring systems have been developed. These systems use either four (SIRS) or six (neonatal SIRS, nSIRS) objective clinical criteria (Supplementary Table 1) to identify foals that potentially have sepsis ^2^. However, these scoring systems have limited sensitivity (SIRS 60%; nSIRS 42%) and specificity (SIRS 69%; nSIRS 76%) for detecting neonatal sepsis ^2^. Bacterial infection, the other hallmark of sepsis, is typically identified through blood cultures, which enable bacteriological identification and subsequent antimicrobial susceptibility profiling. However, the sensitivity of bacterial detection through culture is only 25-45% in foals with sepsis ^16–18^. Quantitative PCR (qPCR) systems have a higher sensitivity (87%) ^19–21^, but are only able to detect a finite set of pathogens, leading to false negative results for pathogens not included in the test. Additionally, false positive results can occur in both culture and qPCR in case of transient bacteremia or sample contamination ^22^. As a result of the low sensitivity and specificity of current diagnostic tools, many newborn foals with sepsis remain undiagnosed or misdiagnosed. Given that foals can deteriorate rapidly within hours, there is an urgent need for improved diagnostic tools for earlier clinical intervention. Thus, expanding diagnostic capabilities and enhancing our understanding of sepsis-causing bacteria in foals is essential.

Cell-free DNA (cfDNA) are short DNA fragments found in body fluids, including plasma, which are released upon cell and microorganism death ^23^. In human medicine, sequencing of plasma microbial cfDNA shows great promise for detecting bacterial pathogens in conditions including sepsis ^12,24,25^. The advantages of cfDNA short-read sequencing include its culture-independent nature, a reasonable turnaround time of 2-3 days (with the potential to speed this to less than one day using alternative sequencing platform ^26^), and the ability to facilitate the unbiased discovery of new pathogens that have not been previously cultured ^24,25^. High-throughput sequencing of cfDNA may also reveal general differences in microbial composition in plasma associated with disease development ^12,24^. The abundance and characteristics such as fragment lengths, fragment end-motifs, and mapping locations of cfDNA molecules, can reveal information on physiological and pathological processes such as the immune response ^27–29^. In human plasma, approximately 99.5% of the cfDNA originates from the host ^30^, which are typically ∼167 bp in length ^31,32^. Microbial cfDNA is shorter, with a substantial fraction being smaller than 100bp in plasma ^33–35^. Taxonomic classification and quantification of the microbial cfDNA provides a multi-pathogen, minimally invasive, accurate assay for diagnosing sepsis in humans ^24,25^, with cfDNA end-motifs potentially enhancing the overall diagnostic process ^36,37^.

In this study, our primary objective is to assess the potential of blood cfDNA sequencing for detecting elevated cfDNA levels of bacteria associated with sepsis in newborn foals with nSIRS. The secondary objectives focus on analyzing the overall cfDNA bacterial composition and investigating host cfDNA factors, including their origin and end-motif. While previous research has focused on cfDNA concentrations in septic foals ^38,39^, sequencing of cfDNA has not been previously conducted, marking this research as a pioneering effort in the field. This endeavor prompted us to create a specialized **f**oal <u>cf</u>DNA sequencing for **b**acterial **i**dentification (<u>cf</u>**FBI**) workflow, incorporating both wetlab and open-sourced computational workflows optimized for detecting pathogenic bacteria in foals with suspected sepsis. By applying the cfFBI pipeline to 32 newborn foals we aim to assess the viability of this approach not only as an alternative diagnostic tool, but also as a method to deepen the understanding of the pathophysiology associated with nSIRS.

## Results

### cfDNA sequencing in newborn foals with sepsis using cfFBI

To investigate the potential of cfDNA sequencing in the context of equine neonatal sepsis, we prospectively included 25 newborn sick foals admitted to Utrecht University Equine Hospital in The Netherlands as well as seven healthy newborn foals (H) (Fig. 1a; Supplementary Table 2, 3). All foals included in this study were between 0 and 6 days of age (Supplementary Tables 2-3). Based on the nSIRS criteria (Supplementary Table 1) ^2^, 11 of these foals were nSIRS-positive (S+; nSIRS≥3), four were nSIRS-negative with zero positive nSIRS parameters (nS-; nSIRS=0), and 10 were nSIRS-negative but had one or two positive nSIRS parameters (sS-; nSIRS=1-2) (Fig. 1a; Supplementary Table 3). Three of the eleven S+ (27%), two of the four nS- (50%) and three of the ten sS- (30%) foals had a positive bacterial blood culture (Supplementary Table 3). We focussed the analysis on comparing S+ against nS- and/or H foals, not sS- foals, as the presence of clinical sepsis-related signs in the sS- group (Supplementary Table 3) suggest that some of these foals could have a bacterial infection or even sepsis. nS- and H foals together represent a realistic clinically relevant background, especially as all nS- samples are derived from the same hospital setting as the S+ samples, ensuring that we account for potential biases related to sample handling and environmental factors.

**Figure 1.**
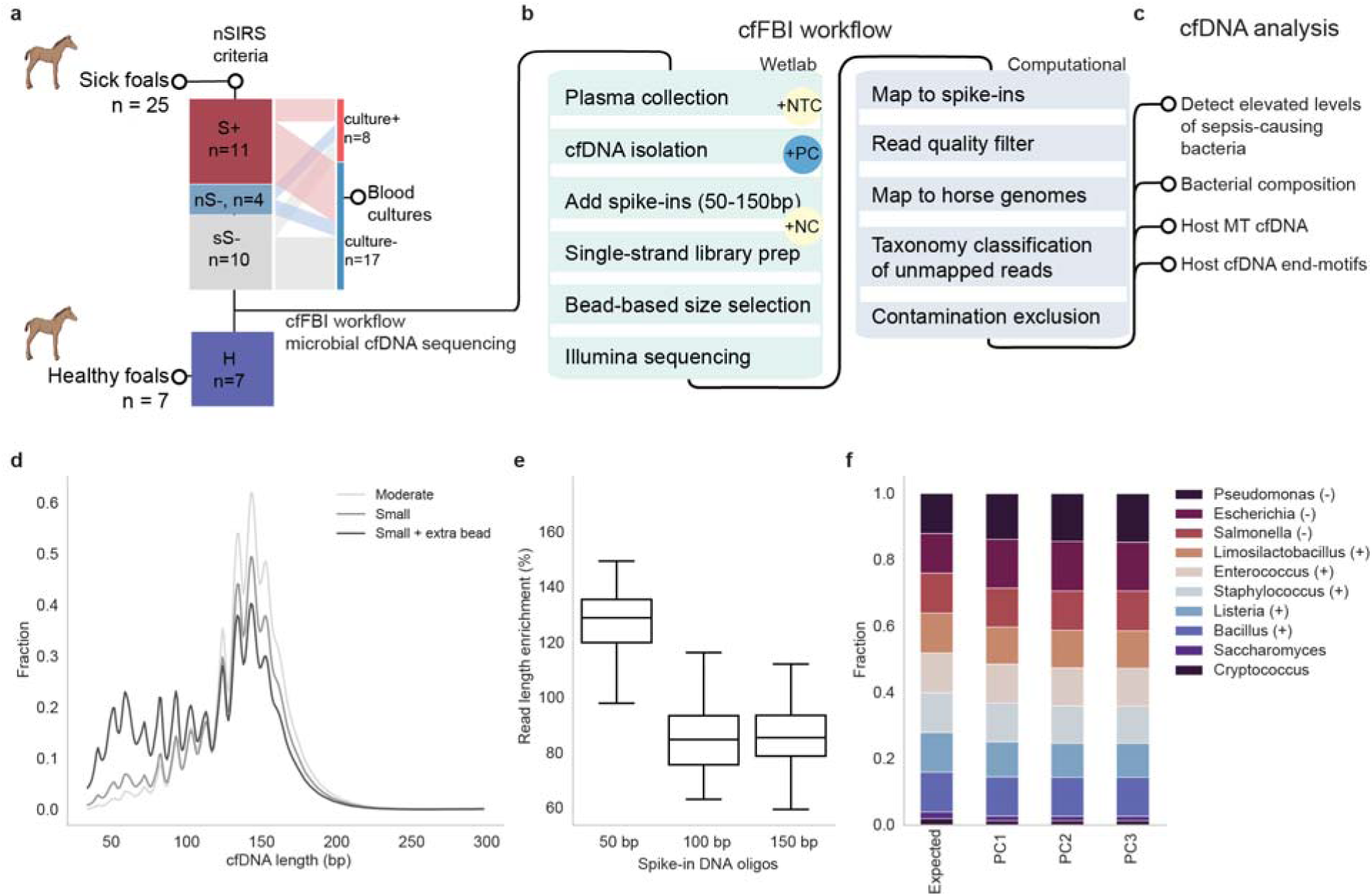
cfFBI pipeline, a cell-free DNA sequencing workflow designed to enhance bacterial identification in foals suspected of sepsis. **a.** Foal cohort and SIRS categorization: The cohort includes sepsis-suspected foals and healthy (H) controls. Foals were categorized based on nSIRS criteria as SIRS-positive (S+; nSIRS ≥ 3), SIRS-negative with no symptoms (nS-; nSIRS=0), or SIRS-negative with symptoms (sS-; nSIRS=1-2). The alluvial plot shows the number of ill foals with positive blood cultures at hospital admission. **b.** Schematic of cfFBI workflow: cfDNA is isolated from foal blood plasma and mixed with synthetic DNA oligos (50, 100, 150 bp). A ligation-based library preparation and bead-based size selection enrich for short microbial fragments (<100 bp). After paired-end Illumina sequencing, spike-in sequences and low-quality reads are filtered out. Remaining reads are mapped to host genomes, and unmapped reads are classified taxonomically using Kraken2 with a customized database. Suspected contaminants are finally excluded. The workflow includes diverse controls: positive controls (PC), no-template controls (NTC), and negative controls (NC). **c.** Comparative analyses were performed within this study, comparing S+ versus H and nS-. Specifically, we focused on variations in host mitochondrial (MT) cfDNA, chromosomal host cfDNA end-motifs, bacterial load and diversity, and the abundance of potential pathogenic bacteria in septic foals. **d.** Comparison of three ligation-based single-strand library preparation methods for enriching short cfDNA fragments: the ’moderate small’ and ’extreme small’ protocols from the SRSLY NGS Library Prep Kit, plus an additional bead-based size selection after the ’extreme small’ protocol. The plot displays the template length size distribution of host cfDNA reads for each method. **e.** Boxplot showing the enrichment or depletion of synthetic DNA oligos (50, 100, and 150 bp) in foal plasma samples (n = 32). Synthetic oligos of 50, 100, and 150 bp were spiked in at equimolar ratios. **f.** Stacked bar plot showing the taxonomic classification results for a sonicated mock community with 10 microbial species. The symbols (-) and (+) after each bacteria genus name indicate whether the species is Gram-negative or Gram-positive. It compares expected versus observed species ratios from three technical replicates using the cfFBI workflow and Bracken abundance re-estimation.

To enable assessment of elevated levels of pathogenic bacteria in the S+ population, we created a cfFBI wetlab and computational workflow (Fig. 1a-c). Given that the microbial cfDNA fraction is known to be minute in plasma ^33^, cfFBI employs a specialized wetlab strategy to enrich for bacterial cfDNA molecules while remaining untargeted in multi-pathogen detection. The cfFBI wetlab cfDNA workflow therefore consists of a ligation-based single-stranded cfDNA library preparation method followed by a bead-based size selection step which effectively enriches short (<100bp) fragments (Fig. 1b-e), which is the known size range for bacterial cfDNA ^40^.

For all foal plasma cfDNA sequencing libraries, between 10 and 50 million paired-end reads were obtained (Supplementary Fig. 1a; Supplementary Table 4). Since bacterial fractions account for less than 0.5% of the total cfDNA even after enrichment ^30^, we deemed it crucial to prevent misclassification,especially false positives. cfFBI’s computational pipeline tackles this challenge through a multi-step process designed to minimize such errors. Previous findings demonstrate a reduction of false positive microbial counts by mapping to a more comprehensive host reference genome ^41^. Therefore, cfFBI maps to the latest horse reference genome, EquCab3, along with all other horse genomes available on NCBI, totaling 11 genomes (Supplementary Fig. 2a,b), increasing the average host fraction of total cfDNA by 0.5%. Second, cfFBI taxonomically classifies the remaining unmapped reads using Kraken2 ^42^, with a custom database that includes all 11 horse genomes, as well as human genomes and all NCBI complete microbial genomes which improves species assignment. Third, suspected microbial contaminants are excluded from downstream analyses by testing whether the levels of microbial species correlated with the volume of reagents used in cfFBI ^43^.

To ascertain the consistency and effectiveness of the cfFBI workflow across library preparations, we first tested the classification accuracy using positive control samples (PCs) from sonicated mock microbial community DNA containing eight bacterial species and two yeasts. In all three technical replicate PCs, the eight bacterial genera were detected at levels consistent with the known microbial composition ^44^, with relative abundance being highly similar across PCs (Fig. 1f; Supplementary Table 5). The consistent detection of the correct microbes in the appropriate ratios across the three different PCs, shows robustness in both the wetlab and the bioinformatics workflow.

### Decontamination and bacterial species composition assessment

We first analyzed the bacterial species composition in the samples, concentrating on both the bacterial fraction and its diversity. Typically 4.2% of the non-mapped reads were confidently classified using Kraken2 (Confidence threshold of 0.8; Supplementary Fig. 2b), of which 1.1% were classified as a bacteria at species level. To ensure accurate analysis of true biological signals, we first removed potential contaminants (see *Methods*), excluding 0.00319% of bacterial reads classified at the species level that were identified as contaminants (Supplementary Table 5). Most of these contaminant species were detected in the negative controls as well (i.e. NTCs and NCs; for details on sample collection see *Methods* and Fig. 1b; for results see Supplementary Table 6 and Supplementary Figs. 3-4), indicating that they are likely contaminants from cfDNA isolation or library preparation. The remaining cfDNA reads classified as bacterial species and not identified as contaminants were aggregated for all downstream bacterial composition analyses.

After decontamination, the median bacterial species-classified cfDNA fraction was 0.0083% (range 0.0010-5.5%) (Fig 2a). This total bacterial fraction moderately correlated with age (r=0.43, Supplementary Fig. 5a,b) and was more variable in the S+ group compared to the other two groups, with some samples showing notably high levels, including one outlier at 5.5% (Fig. 2a,d). Although most of the foals with a high bacterial fraction (above 0.0002) were S+ foals (4/6; 66%), the difference between groups was not statistically significant (Kruskal-Wallis test with Dunn’s multiple comparison tests) (Fig. 2d,e). Between 75 and 1126 (median 327) species were found across 50 to 525 (median 192) genera in each foal. Interestingly, different foals exhibited distinct top abundant species (Fig 2b,c). In total, 4,284 bacterial species across 1,250 different genera were detected, with *Actinobacillus*, *Acinetobacter*, *Streptococcus*, and *Flavobacterium* being the most prevalent, contributing a median of 12.3% per foal.

**Figure 2.**
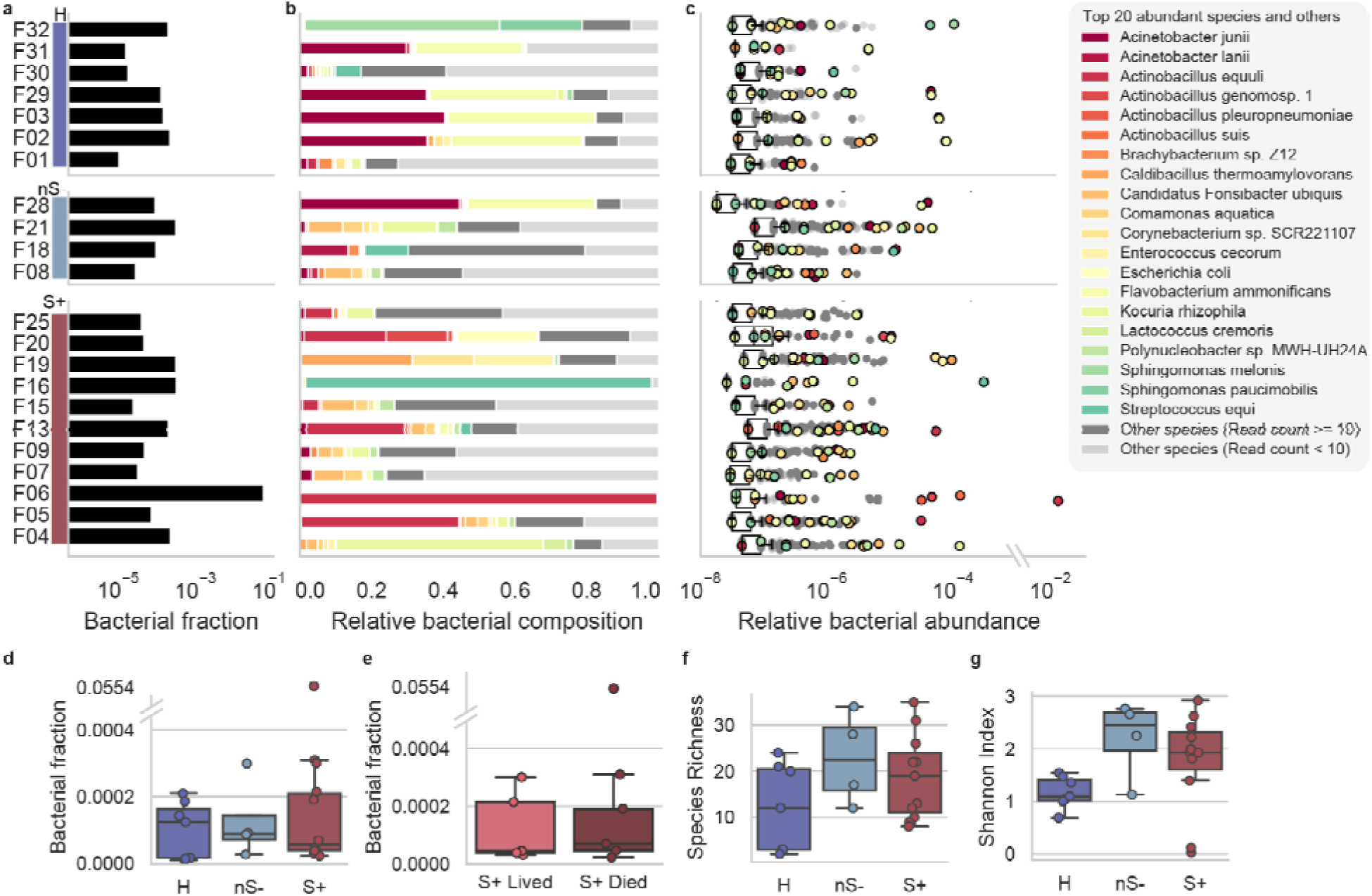
cfDNA bacterial load and diversity in foal plasma samples. **a.** Bacterial fraction in each sample in three categories H, nS- and S+, represented in a log scale. **b.** Relative bacterial composition (normalized to total bacterial reads) in each sample. The top 20 abundant species in all samples were colored (see colors in the legend), the other species with more than 10 exact counts were colored with dark grey color and other species with less than 1 exact counts were colored with light grey color. **c.** Relative abundance (normalized to total cfDNA reads) in each bacterial species, represented in a log scale. The top 20 abundant species in all samples were colored (see colors in the legend), the other species with more than 10 exact counts were colored with dark grey color and other species with less than 10 exact counts were colored with light grey color. **d.** Fraction of cfDNA fragments taxonomically classified as bacterial origin (after removal of contaminant species) and its association with disease status. No significant difference is observed between groups. An outlier at 0.0554 is represented with a broken y-axis. Foals in S+ group showed the largest variation compared to the other two groups. (Standard deviation: H: 0.00008, nS-: 0.00011, S+, 0.0167). **e.** Fraction of cfDNA fragments taxonomically classified as bacterial origin (after removal of contaminant species) and its association with severity of disease in the S+ group. No significant difference is observed between groups. An outlier at 0.0554 is represented with a broken y-axis. **f.** Species richness, representing the number of bacterial species identified (after removal of contaminant species) in each plasma sample, and its association with disease status. **g.** Shannon index, indicating the evenness of classified bacterial species distribution (after removal of contaminant species) within each foal plasma sample, and the association between Shannon index with disease status. **d-g.** Disease status groups are H (n = 7), nS- (n = 4), and S+ (n = 11). For severity of disease, focus is on S+ cases with either survival (n = 5) or death (n = 6). Boxes represent the 25th percentile (bottom), median, and 75th percentile (top), with whiskers extending to the rest of the distribution within 1.5 times the inter-quartile range.

Samples exhibited high variability in species composition (Fig. 2b,c), with the Shannon index revealing greater microbial diversity in sick foals (nS- and S+) compared to healthy foals (Fig. 2f,g), with age having no significant impact (Supplementary Fig. 5c,d). However, neither species richness nor diversity metrics (Supplementary Fig. 6) effectively distinguished between healthy, nS-, and S+ foals (Fig. 2b,c).

### Co-elevation of multiple sepsis-causing genera observed in foals with sepsis

The primary objective of this study is to evaluate the potential of blood cfDNA sequencing for detecting elevated levels of sepsis-causing bacteria in newborn foals with nSIRS. To pinpoint bacteria associated with sepsis in the S+ foals, we compared the aggregated counts of species from the 16 most frequently cultured pathogenic genera found in culture-positive foals with sepsis (Fig. 3; Supplementary Fig. 7-9) ^45^. One or multiple pathogenic genera were higher in 8/11 of the S+ foals compared to both nS- and H foals while the other 3/11 were higher compared to H foals alone (Supplementary Fig. 8), meaning that all S+ foals showed elevated levels of at least one pathogenic genus. In the follow-up analysis, we combined these results for species that showed an increase when compared to either H foals or both nS- and H foals.

**Figure 3.**
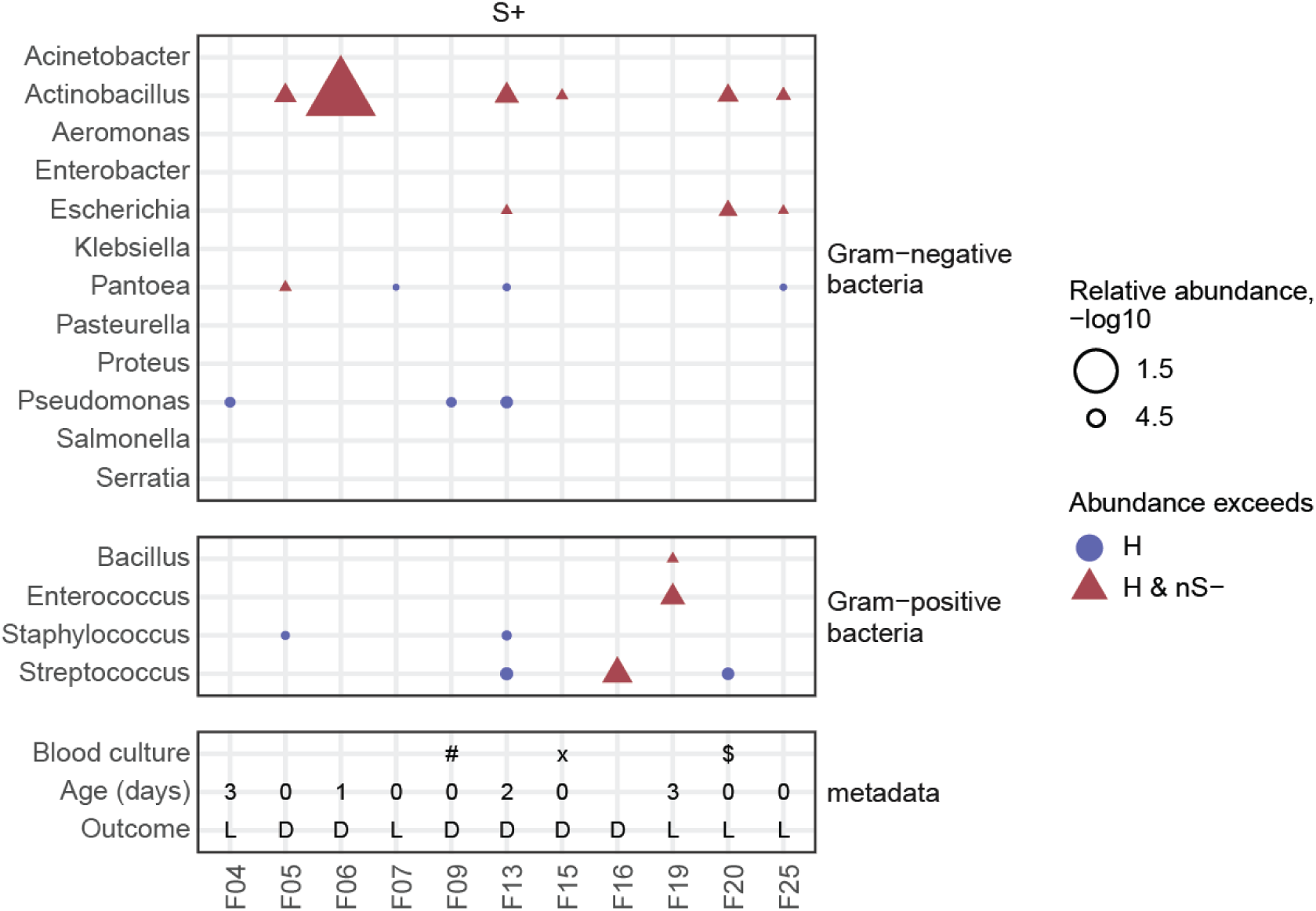
Species abundance and bacterial co-elevation detection in foal cohort samples. Dotplot displaying the detection of the 16 most frequently cultured pathogenic genera. Gram-negative species are shown at the top, and Gram-positive species are shown in the middle. Dots represent genera detected with at least 10 reads. Blue circles indicate genera with a relative abundance higher than in H foals, while red triangles denote genera with a relative abundance higher than in both H and nS- foals. Metadata is represented at the bottom, including blood culture if positive, age at hospital presentation if known, as well as survival outcome (L, Lived; D, Died). Special symbols in the blood culture section: ‘#’ Gram-negative rod (non-fermenter), ‘x’ *Actinobacillus equuli* & *Streptococcus pneumoniae* and ‘$’ *Staphylococcus* coagulase-negative.

Overall, *Actinobacillus* and *Pantoea* were most frequently increased in 6/11 (54.5%) and in 4/11 (36.4%) of the S+ foals, respectively (Fig. 3; Supplementary Fig. 7; Supplementary Table 7). Co-elevation of multiple genera occurred in 5/11 (45.5%) foals (Fig. 3), with co-elevation of *Actinobacillus* and *Escherichia* being most common (3/11; 27.2%; Fig. 3). On the contrary, *Acinetobacter* was the only genus with higher frequencies in multiple H and nS- foals compared to S+ foals (Supplementary Fig. 9), suggesting that the elevated relative abundances of the 16 genera tested represent a genuine biological signal specific to the S+ foals. We did not observe clear relationships between bacterial elevation and the survival outcome of S+ foals (Fig. 3), suggesting that survival may be influenced by factors beyond the bacterial elevation, including the foal’s immune response and the reaction to treatment. Taken together, these microbial cfDNA sequencing results show that sepsis in foals may have a multi-bacterial nature. Furthermore, the results emphasize that microbial cfDNA sequencing may hold potential for newborn foal sepsis diagnosis, although larger studies are required to establish the sensitivity and specificity of the technique.

Given this promise, we further investigated the sS- foals, which were excluded from previous analyses due to the ambiguous disease state. Based on the low sensitivity of the nSIRS criteria (42%) ^2^ and clinical symptoms observed in sS- foals, we expect some foals with sepsis in the sS- group as well. *Acinetobacter* levels were elevated in 4/10 sS- foals compared to S+ (Supplementary Fig. 9), resembling the H foals. Conversely however, the majority of sS- foals displayed trends similar to S+ foals, including increased levels of *Actinobacillus* (6/10) and *Escherichia* (4/10), when compared to H alone or H and nS-. Additionally, 70% of sS- foals exhibited co-elevation of multiple genera (Supplementary Fig. 8). The similarities in bacterial co-elevation between sS- and S+ foals, coupled with the low sensitivity of the nSIRS criteria ^2^, suggest that some foals with sepsis may have been overlooked. Alternatively, it could mean that the foals with a low nSIRS score are in an earlier stage of sepsis-development or suffer from other bacterial infections, both leading to an increase in microbial levels without many clinical nSIRS-symptoms.

Species-level bacterial identification can be used for clinical decision making, including guidance on selection of antimicrobial treatment. Therefore, we evaluated bacterial species-level elevations in the 16 most common genera associated with foal sepsis. Across the 16 genera, 22 pathogenic species were elevated in one or multiple S+ foals (Supplementary Table 8, Supplementary Fig. 7). *Actinobacillus equuli*, *Actinobacillus pleuropneumoniae* and *Escherichia coli* were most frequently elevated in 6/11, 3/11 and 3/11 S+ foals, respectively. Most elevated species corresponded with their respective elevated genera (31/35 observations). However, *Staphylococcus equinus* was higher in F05, *Acinetobacter haemolyticus* was elevated in F04, and *Acinetobacter lanii* as well as *Acinetobacter wanghuae* were elevated in F20, suggesting that species-level information can provide some additional leads compared to genus-level analyses (Supplementary Table 8). However, our analysis also urges caution interpreting sequencing results, particularly about potential misassignment of reads to closely related species, such as the identification of *Actinobacillus pleuropneumoniae*, which is not typically listed as a sepsis-causing species for foals.

Bacterial culture is the golden standard for identifying bacteria ^18,22^, but suffers from low sensitivity and false positive observations ^16–18^. To evaluate the concordance between traditional culture and bacteria identified (as elevated) by microbial cfDNA profiling through sequencing, we compared the results of genus-level blood cultures to cfDNA sequencing after excluding potential contaminants. Notably, elevated levels of sepsis-causing bacterial genera were found by cfDNA sequencing in all three S+ and three sS- foals with positive bacterial blood cultures, indicating that cfDNA sequencing effectively detects bacterial abnormalities associated with sepsis (Supplementary Fig. 9). The overlap between detected genera, however, was limited, with only 43% (3/7) of the culture-identified bacterial genera showing elevation in the cfDNA (Supplementary Table 9). In an additional 29% (2/7) of cases, reads had been assigned to the cultured genera, but cfDNA levels did not surpass those as detected in H and nS- foals, suggesting that the bacteria can be simply present rather than elevated (Supplementary Table 9). Taken together, we observe low concordance between culture and cfDNA-based pathogen identification in newborn foals with SIRS, potentially due to the fact that cfDNA sequencing detects presence and elevated levels of DNA of sepsis-causing bacterial taxa, while culture detects viable bacteria.

Associations of host cfDNA with nSIRS status Since most of the sequenced reads are mapped to the host reference genome and it is recognized that these host-derived reads can offer insights related to infection related tissue damage ^46^, host response to infection ^23^ and sepsis ^47,48^, we next investigated differences in host cfDNA between S+ and H and/or nS- foals, and between S+ foals that lived to S+ foals that died. Confirming previous results in foals ^38^, but differing from observations in humans ^47–49^, total cfDNA levels in plasma were not significantly elevated in S+ foals compared to H and nS- foals (Kruskal-Wallis with Dunn’s multiple comparison; S+ vs. H: p=0.92, Z=0.73; S+ vs. nS- p>0,99, Z=0.31), nor in S+ foals that lived compared to S+ foals that died (Mann-Whitney U Test, p=0.32, U=9, Fig. 4a,b). This suggests that total cfDNA levels cannot be used to diagnose sepsis in foals, as previously reported ^38^. Strikingly, opposite to MT cfDNA levels in human sepsis patients ^49,50^, the MT cfDNA fraction of foal host origin was significantly lower in S+ foals compared to H foals (Fig. 4c). Similarly, a significant decrease in MT cfDNA was observed in S+ foals that died compared to those that survived (Fig. 4d). Of note, none of these variables were significantly different between isolation batches, library preparation batches, and operators (Mann-Whitney U Test with Bonferroni Correction, Supplementary Fig. 10).

**Figure 4.**
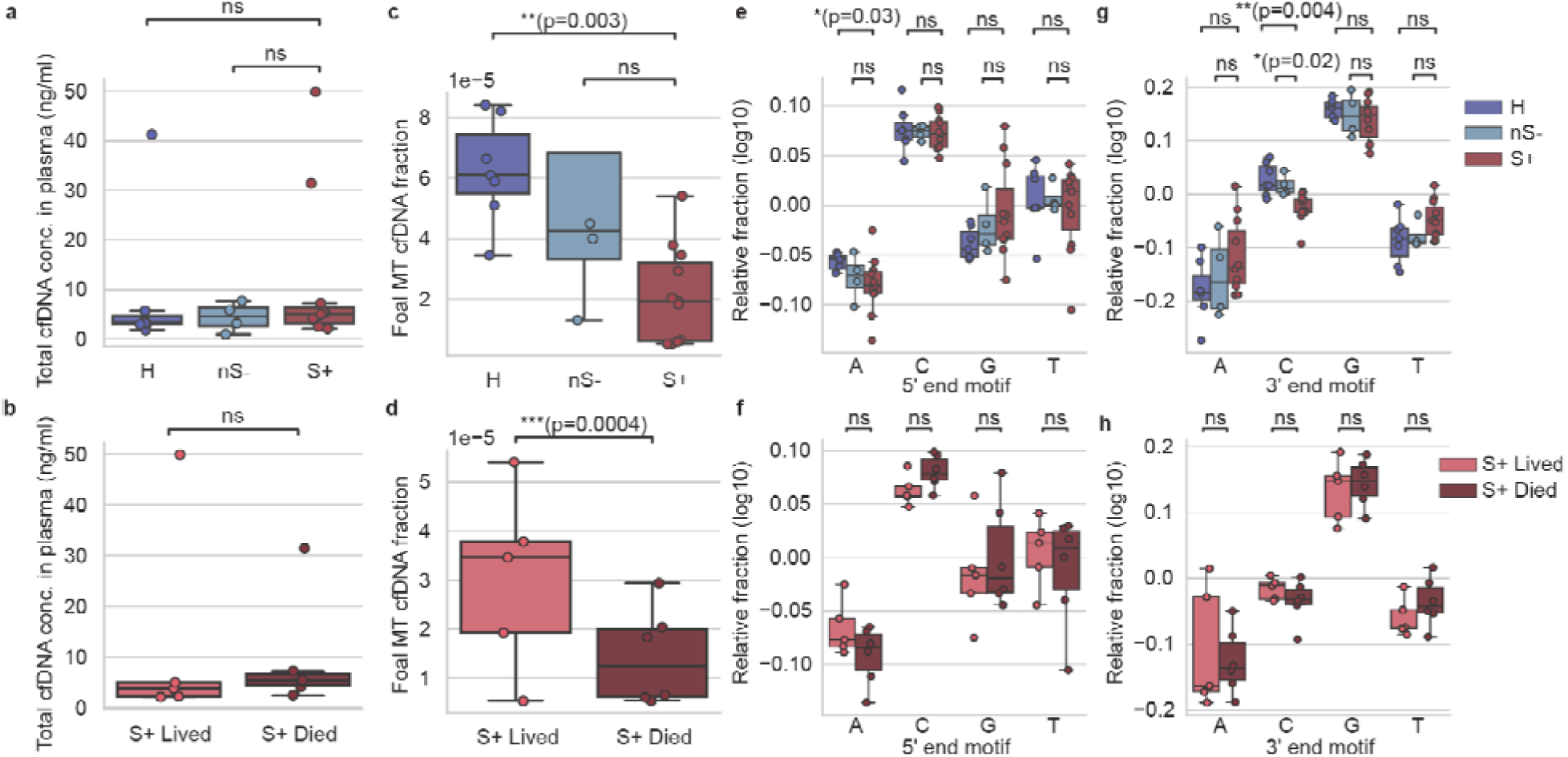
Host cfDNA abundance and its association with disease status and severity of disease. **a.** Total cfDNA concentration in plasma samples and its association with disease status. **b.** Total cfDNA concentration in plasma samples and its association with its severity of disease. **c.** Fraction of cfDNA fragments of host mitochondrial origin, and their association with disease status. **d.** Fraction of cfDNA fragments of host mitochondrial origin, and association with severity of disease. **e.** Normalized base content fraction at the 5’ end of host chromosomal cfDNA fragments and its correlation with disease status. **f.** Normalized base content fraction at the 5’ end of host chromosomal cfDNA fragments and its association with severity of disease. **g.** Normalized base content fraction at the 3’ end of host chromosomal cfDNA fragments and its association with disease status. **h.** Normalized base content fraction at the 3’ end of host chromosomal cfDNA fragments and its association with severity of disease. Disease status groups are H (n = 7), nS- (n = 4), and S+ (n = 11). For severity of disease, focus is on S+ cases with either survival (n = 5) or death (n = 6). Boxes represent the 25th percentile (bottom), median, and 75th percentile (top), with whiskers extending to the rest of the distribution within 1.5 times the interquartile range.

As host end-motifs can give insight into the activities of nucleases and the interplay with innate immune response such as NETs ^28,51^, we proceeded to investigate host chromosomal cfDNA end-motifs. An enrichment in 5’ C-end and 3’ G-end cfDNA reads was present in all samples (Fig. 4e-h, Supplementary Table 10). Specifically 3’ C-end cfDNA reads were significantly decreased in S+ foals compared to H and nS- foals, while 5’ A-end cfDNA reads were significantly decreased in S+ foals compared to H foals (Kruskal-Wallis test with Dunn’s multiple comparison tests, S+ vs. H: p=0.004, Z=3.09; Fig. 4e,g, Supplementary Table 10). Collectively, these results indicate that mitochondrial cfDNA levels and end-motifs could serve as potential biomarkers for SIRS and its prognosis in foals.

## Discussion

We introduce cfFBI, a cell-free DNA sequencing workflow designed to enhance bacterial identification in foals through a combination of optimized wetlab and computational procedures. cfFBI specifically enriches small, microbial cfDNA molecules which are present at minute levels within the cfDNA pool derived from a single tube of blood. cfFBI’s computational steps, including host mapping and decontamination, are optimized to minimize false bacterial identifications. Using cfFBI, we applied cfDNA sequencing to newborn foals for the first time and detected elevated bacterial levels in 8/11 nSIRS-positive foals compared to levels in (sick) nSIRS-negative foals, while the other 3/11 nSIRS-positive foals showed at least one bacterial genus elevated compared to healthy foals alone. Interestingly, we find co-elevation of multiple pathogenic bacteria in 5/11 (45.5%) of nSIRS-positive foals. Although it was already known that bacterial culture can provide positive results for multiple species ^11–14,52–54^, it remained unclear if these were true co-occurrences, similar to the co-occurrence observed in human sepsis patients ^12,13^ or a result of contamination ^17,18,55^. The frequent observation of co-elevation in cfDNA in this study suggests that multiple genera may actually jointly contribute to sepsis in newborn foals. Further validation and follow-up research is needed to determine the potential implications of these findings.

Until now, most knowledge about the bacteria causing sepsis in foals has been based on culture-dependent techniques, while culture is known to have only 25-45% sensitivity in foals with sepsis ^16–18^. When comparing bacteria detected by cfDNA sequencing to those identified through culture in the cohort, we observed limited concordance (3/7 (43%)). This discrepancy may arise because bacterial culture detects only viable bacteria, while cfDNA sequencing reveals both the presence and increased abundance of bacterial cfDNA resulting from recent cell death. The two techniques thereby capture a different aspect of the complex underlying pathophysiology. In large-scale human studies, microbial cfDNA showed higher sensitivity and specificity than blood cultures for detecting clinically relevant pathogens, resulting in an enhanced patient survival, and a reduction of overall antimicrobial use in patients with sepsis ^56,57^. Ultimately, the two techniques may turn out to be complementary, with cfDNA sequencing providing reliable multi-pathogen results, while culture provides valuable insights into antimicrobial resistance of the identified bacterial species.

Diagnosis of sepsis in newborn foals is challenging, with most tools suffering from a low sensitivity and specificity. This issue also applies to the nSIRS scoring system used here, where foals with sepsis can have a nSIRS score of less than 3 ^2^. To minimize the impact of potential false negative septic foals in the nSIRS-negative group, we excluded the nSIRS-foals with a nSIRS score of 1-2 when setting background level for bacterial elevation analysis in nSIRS-positive foals, as some of these cases may have an unindicated sepsis or a bacterial infection. Simultaneously, the nSIRS-positive group may still include foals without sepsis, so without a bacterial infection, as sepsis is currently defined as a combination of SIRS with a bacterial infection. This challenge clearly shows the need for additional diagnostic tools for improved sepsis diagnosis in foals.

Although this study clearly indicates the potential of cfDNA sequencing in newborn foals for the first time, it should be noted that the current cohort has limited statistical power to detect significantly elevated microbes in nSIRS-positive foals above background. We opted for a conservative approach of identifying elevated bacterial levels by testing if the value is above the highest observation in the control background samples without aiming to test for significance. A larger cohort study involving more newborn foals is essential to fully assess the potential of cfDNA sequencing for diagnosing sepsis. This would also allow a comparison to healthy and nSIRS-negative foals separately, where the first comparison is informative to gain more insight into biology, whereas the latter comparison can provide tools useful in the clinic. Increased cohort size would also be beneficial to study abundance of bacterial species or genera in a more unbiased manner, without focusing exclusively on the 16 most frequently observed bacteria in culture. This could provide new insights into biologically relevant, yet hard-to-culture, bacterial taxa associated with sepsis.

In addition to validating the microbial cfDNA observations of the current study, a larger study could aim to further explore the trends we found in the host cfDNA. This includes confirming the decreased MT cfDNA fraction in nSIRS-positive foals which aligns with previous studies in foals ^38,39^, but contrary to what is observed in humans ^49,50^. Moreover, such a study would enable the validation of the absence of the expected elevation in total cfDNA in nSIRS-positive cases ^38,39^, which would typically be indicative of increased tissue damage and cell death as is seen in humans ^58^. It could also shed light on findings on the (complementary or concordant) relationship between host cfDNA and microbial cfDNA which we did not observe in the current cohort (Supplementary Table 10). Finally, the differences in end-motifs in host cfDNA in nSIRS-positive versus nSIRS-negative foals could be validated in a larger cohort, potentially revealing additional biomarkers. By combining host and pathogen information from plasma cfDNA, more insights of host transcription profiles such as innate immune response activities can be obtained, as was already shown in human sepsis patients ^23,59^.

Currently, foals suspected of having sepsis are treated with broad spectrum antimicrobials until culture and susceptibility testing results become available after approximately 72 hours. In cases that fail to improve within this time period, antimicrobial treatment regimens are adjusted based on historical information on prevalence and susceptibility of bacteria causing sepsis in foals in that specific geographic area. As sepsis and organ dysfunction can develop rapidly, the antimicrobial susceptibility tests are rarely timely to aid the treatment regimen. By utilizing a faster and more sensitive technique ^26^ for identifying potential pathogens, coupled with the continually decreasing costs of sequencing combined with more targeted approaches (e.g. for antimicrobial resistance genes ^60^), future adjustments to antimicrobial therapy can be made earlier, potentially increasing the survival chances of foals with sepsis.

## Methods

### Foal cohort

We prospectively included 25 sick foals admitted to the Utrecht University Equine Hospital (Utrecht, The Netherlands), between March 1st, 2021 and July 1st, 2022. For diagnostic purposes, two blood samples (up to 20 mL each) and one blood sample of 10 mL were collected aseptically from the jugular or cephalic vein, either by venepuncture or through a newly placed intravenous catheter immediately upon hospitalization. The two 20 mL samples were placed into 70 mL brain heart infusion broth + SPS (Biotrading, Mijdrecht, the Netherlands) and transported to the Veterinary Microbiological Diagnostic Center where the bottles were incubated at 37°C for 18-24h. After incubation, Gram-staining was performed followed by inoculation on two sheep blood agars (SBA), chocolate agar (CHOC) and MacConkey agar (MAC; Biotrading, Mijdrecht, the Netherlands). One SBA and MAC agar were incubated aerobically, while the other SBA was incubated anaerobically and the CHOC agar was incubated microaerobically; all at 37°C for 5-7 days. Agars and broths were checked daily for bacterial growth. If bacterial growth was detected, identification took place using Maldi-TOF (Bruker, Bremen, Germany). The 10 mL sample of blood was collected directly into a Streck tube (see “Sample preparation and nucleic acid isolation“). Diagnostic and clinical data of sick foals were recorded and later extracted from the medical information system (Supplementary Table 2).

In addition to the sick foals, seven healthy (H) newborn foals were enrolled in this study, four from Utrecht University Equine Hospital (Utrecht, The Netherlands) and three from Dierenkliniek Emmeloord (Emmeloord, The Netherlands). In the healthy foals, at the moment of blood collection for the routine check for passive transfer of immunity, 10 mL of blood was collected aseptically for cfDNA sequencing. Blood cultures were not performed on the healthy foal samples.

nSIRS criteria were used to classify foals into groups (Supplementary Table 1) ^2^, whereby an nSIRS score ≥ 3 was considered nSIRS-positive (S+), a nSIRS score of 0 was considered nSIRS-negative (nS-) and an nSIRS score of 1-2 was considered symptomatic nSIRS-negative (sS-). The foal cohort thus comprised 11 S+ foals, four nS- foals, seven healthy foals, and 10 sS- foals. The demographics, including foal age at presentation, breed, sex, as well as dam age, gestation length, and parity, are detailed in Supplementary Tables 2-3.

### Sample preparation and nucleic acid isolation

Blood for cfDNA sequencing was collected aseptically in Streck Cell-Free BCT (Streck #230257). Plasma extraction involved centrifugation for 10 minutes at 1600 g (at room temperature), followed by an additional centrifugation step for 10 minutes at 16,000 g (at 4 °C) to eliminate all cells and debris. The resulting plasma samples were then stored at -80 °C.

For nucleic acid isolation from plasma, the Circulating Nucleic Acid Kit (Qiagen, 55114) was employed with specific modifications to the manufacturer’s protocol. First, a subset of the samples was supplemented to 5mL using PBS (Supplementary Table 4), prior to isolation. Second, the lysis time was extended from 30 to 60 minutes. Finally, cfDNA was eluted in 28 or 35 µL of Nuclease Free water (Invitrogen, 10977-035), and measured by the Qubit dsDNA High Sensitivity Assay Kit or Broad Range Assay Kit (Thermofisher Scientific, Q32854 and Q32853, respectively).

### Sequencing library preparation using single-strand ligation based DNA-capture

For library preparation quality control purposes, plasma DNA was supplemented with synthetic spike-ins, equalling 0.2% of the total DNA input. The synthetic spike-ins consisted of an equimolar mix of three single-stranded DNA sequences that were 50, 100, and 150 bp in length (sequences of these spike-ins listed in Supplementary Table 11). Since the SRSLY splint adapter (refer to the next section for more details about SRSLY) contains a 7-base random overhang, the spike-ins were designed to include a random overhang sequence of the same length.

Then, the SRSLY PicoPlus NGS Library Prep Kit was used to prepare sequencing libraries (Claret BioScience, CBS-K250B-96). Briefly, DNA input molecules were denatured and kept as single-stranded molecules using a thermostable single-stranded DNA binding protein. The single-stranded DNA was then ligated to SRSLY splint adapters, followed by an indexing PCR ^61^. To enrich short fragments in all foal samples, we used the small fragment retention version of the SRSLY PicoPlus NGS Library Prep Kit protocol along with an additional bead-based size selection step (Ampure XP, A63882). In a separate experiment (Fig. 1d) we compared a short fragment retention and moderate fragment retention protocol combined with/without a customized extra step of bead-based selection in a separate experiment, by which we established that we would use the small fragment retention version with a customized extra step of bead-based selection in all other samples in this study. The complete description of this experiment can be found in the method section “Short fragmentation length enrichment analysis”.

All libraries were quantified using the Qubit dsDNA High Sensitivity Assay Kit (Thermofisher Scientific, 32854) and size distribution was analyzed using the Tapestation 2200 and the D1000 kits (Agilent, 5067-5583). Foal sample sequencing libraries were pooled equimolar, with positive (see “Preparing positive control samples imitating microbial cfDNA fragments”) and negative controls (see “Negative controls for identifying contaminants in low microbial load samples”), albeit at a threefold lower molar ratio than the foal cfDNA libraries.This pool was subsequently enriched for sub-100 bp cfDNA molecules by a bead-based size selection step (Ampure XP, A63882). After this bead-based size selection, the concentration and size of the library pool were measured using the TapeStation 2200 and the D1000 kit.

### Preparing positive control samples imitating microbial cfDNA fragments

As a positive control, we made use of a sonicated mock community DNA (ZymoBIOMICS Microbial Community DNA Standard, D6305) containing a mixture of genomic DNA of 10 microbial strains: *Listeria monocytogenes, Pseudomonas aeruginosa, Escherichia coli, Salmonella enterica, Lactobacillus fermentum, Enterococcus faecalis, Staphylococcus aureus, Bacillus subtilis, Saccharomyces cerevisiae* and *Cryptococcus neoformans*. In short, 2 µL ZymoBIOMICS standard was supplemented with 88 µl of LowTE (10mM Tris, 0.1mM EDTA), before shearing using the Covaris S2 at 6-8°C, with continuous degassing, a duty cycle of 10%, intensity set to 5, and 200 cycles per burst for 14 minutes. Bead-based size selection was then performed to enrich for DNA fragments shorter than 200 bp (Ampure XP, A63882), using an initial 1.1x volume of beads followed by adding a 3x volume of beads to the supernatant, to mimic cfDNA. Three ng of sheared, size-selected mock community DNA supplemented with 6 pg of synthetic spike-in DNA were used as input for the SRSLY library preparation. Since the next-generation sequencing libraries were prepared in three separate batches, we included one positive control sample for each batch, resulting in a total of three positive controls (PC1, PC2, and PC3).

After sequencing, the computational cfFBI workflow was applied to the positive control (PC) samples, including Bracken abundance re-estimation to refine the relative fractions of each species and genus (see “Sequencing Data Processing Using the cfFBI Pipeline” for details). The observations for these 10 species and their respective genera are provided in this study, including their relative fractions at both the species and genus levels, as well as the variance among the PC samples.

Of note: the positive controls (PC1-PC3) served three purposes. First, to validate the effectiveness of the protocol in each experimental batch. Second, to ensure the wet lab and computational workflow can accurately produce representative species and genera of interest. Third, to confirm that data generated from independent library preparations are comparable.

### Negative controls for identifying contaminants in low microbial load samples

Due to the risk of contamination in low microbial load samples, we incorporated a set of four negative controls (NTC1, NTC2, NCMQiso, NCMQlib). Among these, two (NTC1, NTC2) consisted of 5 ml PBS that underwent the entire process of cfDNA and SRSLY-mediated NGS sequencing library preparation. Another two control samples contained Nuclease-Free water that was utilized for the elution (NC1MQiso) of cfDNA after cfDNA isolation and the supplemention of up to 18 ul that was added to samples before library preparation (NC1MQlib). These two samples underwent the process of SRSLY library preparation. No spike-in DNA sequences were added to the negative control samples.

### Next-generation sequencing

Library sequencing was executed on the NovaSeq 6000 platform with 2 x 150 bp reads. This process yielded a range of 20 to 66 million reads per cfDNA library, between 8.6 and 10.3 million reads for each positive control (PC), and between 6.0 and 9.1 million reads for negative controls (NTC1, NTC2, NC1MQiso, and NC1MQlib).

### Sequencing data processing using the cfFBI-pipeline

Illumina sequencing and synthetic data underwent processing via the cfFBI-pipeline, available on our Github repository (https://github.com/AEWesdorp/cfFBI/tree/main/pipeline). In a nutshell, bbduk.sh from tool BBmap ^62^ was employed to detect and eliminate reads containing 50mer, 100mer, or 150mer synthetic spike-in sequences. Subsequently, duplicate removal was carried out using nubeam-dedup ^63^, followed by default read quality filtering using fastp^64^ to generate high-quality sequencing data. The quality filtering included removing low-quality reads, implementing a low complexity filter, adapter removal, and discarding short reads (< 35bp) using AdapterRemoval ^65^.

For horse read sequence identification, we tested two strategies via host genome mapping (bowtie2 ^66^). The first strategy utilized the reference genome *Equus caballus* EquCab3.0 from NCBI RefSeq (accessed on Nov 8th, 2022). The second strategy incorporated all 10 additional genomic sequences available for *Equus caballus* within NCBI RefSeq (accessed on Feb 5th, 2024), bringing the total to 11 genomes: EquCab3.0, 57H, 25H, 16H, 7H, 2H, 9H, 30H, Ajinai1.0, LipY764, and EquCab2.0.

The latter strategy is adopted in the cfFBI pipeline. After host sequence subtraction, remaining paired-end reads underwent taxonomic classification using Kraken2 ^42^, a highly regarded metagenomic tool that performs exact *k*-mer alignment to a reference database for rapid per-read taxonomic classification (for details about the adapted database, see: “Taxonomic database construction and taxonomy classification”). Sequencing data were processed with a confidence threshold (CT) of 0.8 for all described databases, the selection of the CT was based on previous work ^67^ which demonstrated that a CT of 0.8 results in the highest average precision when using the NCBI database. After Kraken2 classification, Bracken ^44,67^ was employed to re-estimate the abundance of species within the metagenomic PCs (PC1-PC3; as specified in the cfFBI’s config file). Of note, Bracken abundance re-estimation was applied for PCs but not for foal cfDNA samples according to the Kraken software suite authors’ recommendation ^68^.Resulting host-mapping reads, classified reads, and bacterial-classified reads were normalized to QC-passed reads in each sample unless otherwise specified.

### Taxonomic database construction and taxonomy classification

For this study, we constructed a custom Kraken2 hash-table database that includes 11 horse genomes, two human genomes, and all complete microbial genomes from NCBI (downloaded as of May 15th, 2023). The microbial component comprises 285,825 bacterial, 14,977 viral, 496 fungal, 1,493 archaeal, and 96 protozoal genome assemblies. To build this database, genomic sequences from the NCBI RefSeq database were downloaded using the *kraken2-build --download-taxonomy* command for archaea, bacteria, fungi, human, plasmid, protozoa, UniVec_Core (contaminant sequences), and viral genomes. Additionally, all 11 genomic sequences for Equus caballus (EquCab3.0, 57H, 25H, 16H, 7H, 2H, 9H, 30H, Ajinai1.0, LipY764, and EquCab2.0) from NCBI RefSeq were downloaded (as of 05-02-2024). The database also incorporated the human genome GRCh38.p14 (obtained directly from NCBI RefSeq) and CHM13v2.0 (added manually). Kraken2 databases were built using the default settings (*kraken2-build*).

### Short fragmentation length enrichment analysis

To evaluate the efficiency of short (<100 bp) microbial cfDNA fragment enrichment across various protocols, we tested different versions of the SRSLY PicoPlus NGS Library Prep Kit (Claret BioScience, CBS-K250B-96), including short fragment retention and moderate fragment retention protocol combined with or without a customized extra step of bead-based selection. After preparing these four different libraries, the libraries were pooled and sequenced with NextSeq 2000. On average 29 million paired-end (2 x 150 bp) reads were obtained. The cfFBI computational workflow was applied to all four samples, using only the EquCab3.0 as a reference genome. Reads mapped to the host chromosomal contigs were extracted using samtools (v1.3.1) with a minimal mapping quality score of >= 30. Subsequent processing and length analysis were performed by a customized processing script using R (v4.2.1) (for details, see: https://github.com/AEWesdorp/cfFBI/tree/main/fragmentomics).

### Host mitochondrial and host read end-motifs analysis

Sequences mapped to mitochondrial contig (RefSeq contig name NC_001640.1) and chromosomal contigs of *Equus caballus* EquCab3.0 were extracted for host read mitochondrial and host read end-motif analyses using samtools (v1.19, v1.3.1) . Amount of reads mapped to mitochondria with a quality score of >= 40 was divided by the number of reads mapped to all chromosomal and mitochondrial DNA with a quality score of >= 40 in EquCab3.0 to calculate MT cfDNA fraction. To investigate read end-motifs, we analyzed the most terminal base of R1 and R2 in all reads that were mapped with a quality score of >= 30, using a custom R script. Counts of each terminal base were tallied, then normalized to the expected fraction (equally distributed). As an example, for Motif A in 1-mer end-motif: Motif A Relative fraction (log10) = log10( motifA / (∑(motifA+motifB+motifC+motifD) /4) )

### Contamination identification in sequencing runs with concentration-based methods

Low biomass microbial sequencing is sensitive to any DNA sequence present in samples, including contaminants. Decontamination, which refers to the process of removing contaminants from findings, is crucial to exclude uninformative findings. Previously, “*decontam*”, a statistical method which identifies and removes reagent-related contaminant sequences has been proposed for metagenomics data. This method, implemented in an R package, detects contaminants by analyzing their correlation with DNA concentration (frequency-based method) as well as their presence in negative controls (prevalence-based method) ^43^ (illustration adapted in Supplementary Fig. 4b). We adapted the frequency-based method to identify contaminants resulting from the cfDNA isolation step and/or the SRSLY library preparation step (Supplementary Fig. 4a).

This frequency-based method assumes that contaminants exist at the same concentration in input reagents in different samples. Therefore, contaminants are more abundant in samples with low DNA input and consequently samples with lower DNA yield. We measured the cfDNA yield after isolation and used (whenever possible) 5 ng of cfDNA as input for the library preparation, to standardize the input DNA for this step. As a result, more isolation-related contaminants were expected in lower input DNA concentration samples. After SRSLY sequencing library preparation we again measured the total DNA yield, and pooled samples in equimolar amounts for sequencing, thus, more library preparation-related contaminants were expected in samples with low total DNA yield.

Therefore, both cfDNA isolation yield and SRSLY sequencing library preparation yield were used for contaminant identification. Normalized species classified counts of all species were correlated with (1) the inverse input DNA volume used for the library preparation, against correlation to a constant value; and correlated with (2) final DNA yield after library preparation, against correlation to a constant value. Testing the null hypothesis of whether each species was not a contaminant from step (1) and/or step (2) with the R package *decontam*. This derived a p-value (significance) of the likelihood of being a contaminant for each species. The authors of the decontam package suggested identifying the trough in the distribution of p-values to set as a cutoff to identify contaminants. We set a p-value cutoff at 0.25 for both steps and checked whether the species occurred in at least six samples, as a lenient approach to identify as many suspected contaminants as possible to prevent false positive findings as suspected pathogens (Supplementary Fig. 4f,g). We performed this across all isolation and library preparation samples without batch-specific testing due to small batch sizes in this experiment. The identified contaminant species were removed from the further analysis that involves classification results. To validate that the identified species were true contaminants, we checked their presence in four negative controls (NTC1, NTC2, NC1MQiso, and NC1MQlib, see Supplementary Fig. 5).

### Bacterial load calculation

After excluding contaminant species, read counts of all other bacterial species were aggregated and normalized to the total number of quality-filtered reads, to calculate what we call the “total fraction of bacterial cfDNA”.

### Bacterial diversity measurements

For bacterial diversity measurement, it is important to avoid analytical noise arising from potential false positive taxonomy classification. Apart from removing species deemed as contaminants from the above mentioned method, we also exclude all observations that were fewer than ten classified reads. Species richness was calculated as the number of species present in each sample. This includes only species that have more than ten classified reads in any sample and are not identified as contaminant species. The Shannon index, or Shannon entropy, measures biodiversity by considering both the abundance and evenness of species present in a sample ^69^. We also measured Bray-Curtis dissimilarity to assess compositional differences between sample pairs. In order to calculate this Bray-Curtis dissimilarity for all pairs, counts were log10-transformed. Results were visualized using hierarchical clustering with single linkage (nearest neighbor) to illustrate the relationships between samples.

### Identification elevated bacterial taxa

Using the cfFBI workflow, we aimed to detect elevated levels of frequently observed bacterial pathogens in newborn foals with sepsis ^45^. These bacterial pathogens include 12 Gram-negative genera (*Serratia*, *Salmonella*, *Pseudomonas*, *Proteus*, *Pasteurella*, *Pantoea*, *Klebsiella*, *Escherichia*, *Enterobacter*, *Aeromonas*, *Actinobacillus*, and *Acinetobacter*) and four Gram-positive genera (*Streptococcus*, *Staphylococcus*, *Enterococcus*, and *Bacillus*). We visualized the taxonomy-classified normalized read counts for all species within these 16 genera (Supplementary Fig. 7) and aggregated these counts to form a total count for each genus. Comparisons were made by contrasting the foals’ (e.g., S+) aggregated genus-level counts against the maximum aggregated genus-level counts observed in H and/or nS- foals to identify abundances that exceeded those in the control group (i.e., H and/or nS-). Conversely, we also compared the aggregated genus-level counts of the non-S+ foals against those observed in the S+ group. In all comparisons, genera detected at low abundance (fewer than 10 reads) were excluded.

### Statistics

We utilized Kruskal-Wallis tests followed by Dunn’s multiple comparison tests to conduct a directional non-parametric ANOVA for comparing total cfDNA levels, mitochondrial cfDNA fraction, host cfDNA end-motifs, bacterial cfDNA fraction, species richness, and Shannon indices among groups composed of H, nS- or S+ foals. For the comparison between two groups (S+ Lived and S+ Died) we used Mann-Whitney U tests. Additionally, Mann-Whitney U tests with Bonferroni correction were employed to assess whether variables of interest were confounded by factors such as the isolation batch, library preparation batch, and the location of the sample during library preparation.

### Software

Data and statistical analyses were carried out using Python (v3.10) with the packages numpy (v1.26.0) and statannotations (v0.6.0), and GraphPad Prism (v10.3.0). Figures were created using R (v.4.2.0) and Python (v3.10) with the packages seaborn (v0.11.2), pandas (v1.5.3), and compiled with Adobe illustrator (2024, v28.6). Illustrations were created using BioRender and Adobe illustrator (2024, v28.6).

## Code availability

The code related to analysis and visualization of content in this manuscript are deposited at a Github repository: https://github.com/AEWesdorp/cfFBI. The repository is open access with a GNU general public license version 3.

## Data availability

Metagenomic sequencing data (FASTQ files) have been deposited in the European Nucleotide Archive (ENA) Browser under accession PRJEB77374.

## Acknowledgments

This study is supported by the funding of Stichting Vrienden Diergeneeskunde, the Faculty of Veterinary Medicine, Utrecht University, and a Vidi Fellowship (639.072.715) to JdR from the Dutch Organization for Scientific Research (Nederlandse Organisatie voor Wetenschappelijk Onderzoek, NWO). We thank Rhoxane Korthals from Dierenkliniek Emmeloord for her assistance in sample collection. We thank Joost van Rosmalen for his valuable input on statistical analysis. We acknowledge the Utrecht Sequencing Facility (USEQ) for providing sequencing services and data. USEQ is subsidized by the University Medical Center Utrecht and The Netherlands X-omics Initiative (NWO project 184.034.019). We thank Marc Pagès-Gallego and Dieter Stoker for proofreading the manuscript.

## Author Contributions

LC, EW, MJ, ES, MT, CV, JdR conceptually designed the study. ES and MT collected samples, gathered clinical information, and gave clinical input. EW, ES, NB, CV, MT prepared samples. EW, NB and CV optimized protocols and generated the sequencing libraries. LC, EW, MJ contributed to data analysis. CV, AZ, EB, JW provided input on the experiments and analyses. LC, EW, MJ wrote the manuscript. JdR coordinated the study. All authors read and approved the final manuscript.

## Declaration of generative AI and AI-assisted technologies in the writing process

During the preparation of this work the author(s) used chatGPT in order to rephrase sentences. After using this tool/service, the author(s) reviewed and edited the content as needed and take(s) full responsibility for the content of the publication.

## Declaration of interests

JdR is founder and shareholder of Cyclomics BV, a genomics company. The other authors declare no competing interests.

## Supplementary Figures

**Supplementary Figure 1.**
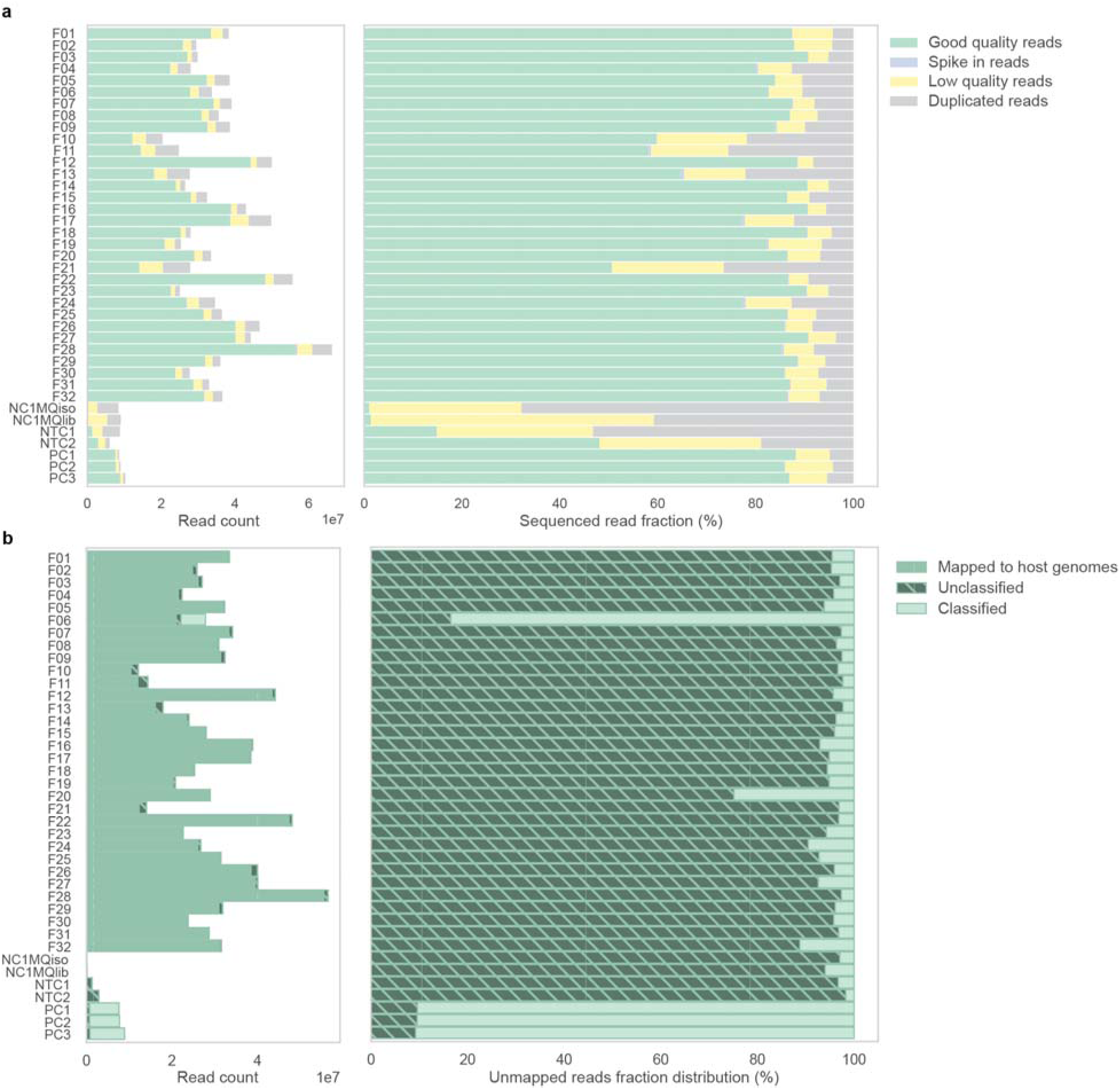
Read counts and fractions per Illumina sequencing library, as determined by cfFBI. **a.** The stacked bar graphs display the number (left) and proportion (right) of paired-end Illumina sequenced reads for each sequencing library. Colors represent the fractions of spike-in reads, duplicated reads, low-quality reads, and high-quality reads, as subsequently determined by the cfFBI workflow pipeline. **b.** The stacked bar graphs illustrate the number of high-quality reads. for each sequencing library. Colors indicate the fractions of reads mapped to the host genome, taxonomically classified and unclassified reads, as subsequently determined by the cfFBI workflow pipeline. **c.** The relative proportion of reads not mapped to host for each sequencing library. Colors the fractions of taxonomically classified and unclassified reads, as subsequently determined by the cfFBI workflow pipeline.

**Supplementary Figure 2.**
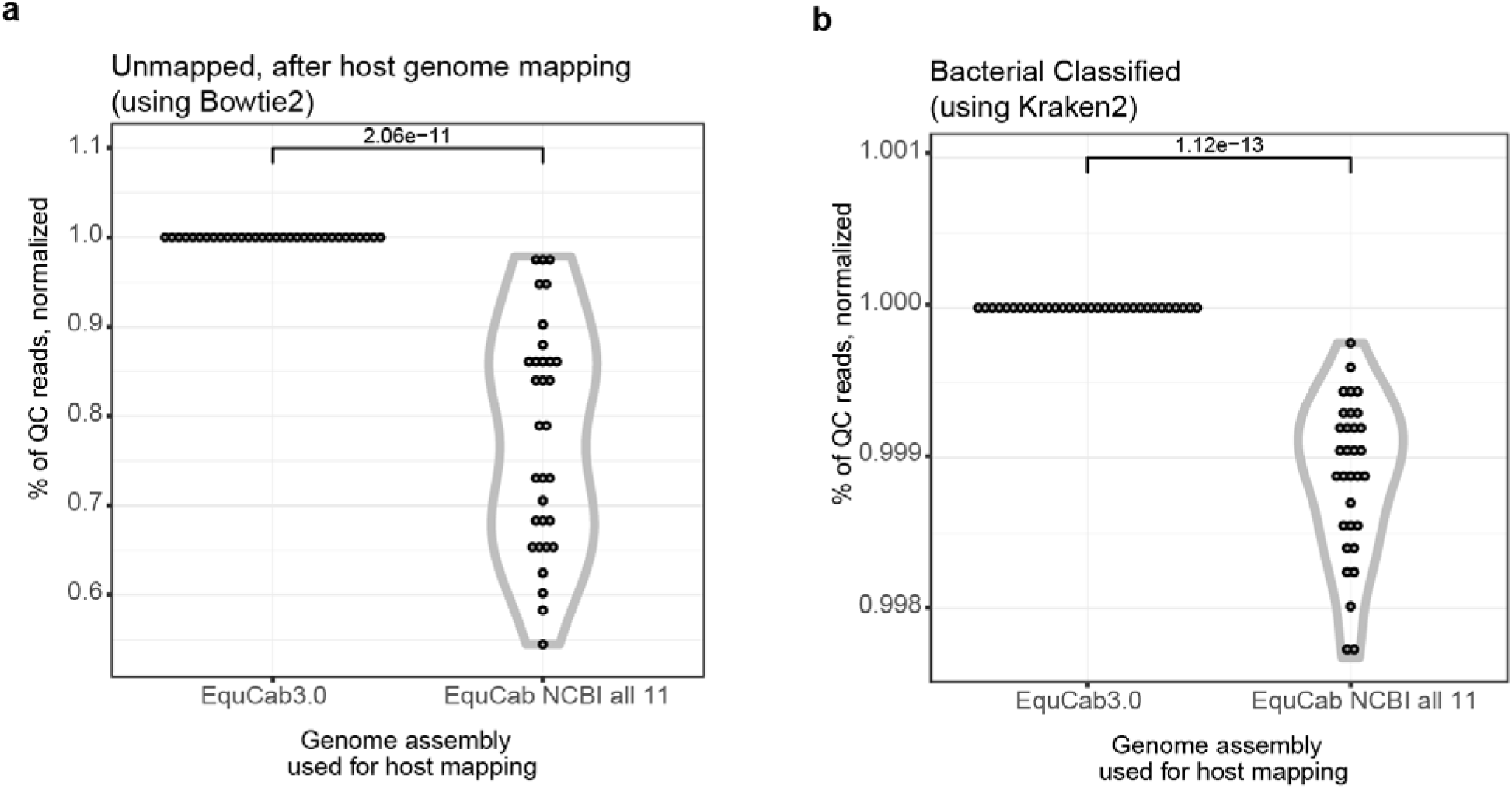
Comparison of read host mapping and bacterial classification using different horse genome reference genomes. **a.** Percentage of quality-controlled reads remaining after mapping to the human reference genome using the cfFBI workflow. Two conditions were compared: using the EquCab3 reference genome alone, and using a compendium that includes all available horse genomes on NCBI (a total of 11 genomes; EquCab NCBI all 11). The results are normalized to those obtained with EquCab3. Statistical analysis was conducted using one-tailed paired t-tests after normalization. **b.** Percentage of quality-controlled reads classified as bacterial by Kraken2, after subtracting host reads via reference genome mapping. Two conditions were compared: using the EquCab3 reference genome alone, and using a compendium that includes all available horse genomes on NCBI (a total of 11 genomes; EquCab NCBI all 11). The Kraken2 bacterial classification results are normalized to those obtained with the EquCab3 reference genome. Statistical analysis was conducted using one-tailed paired t-tests after normalization.

**Supplementary Figure 3.**
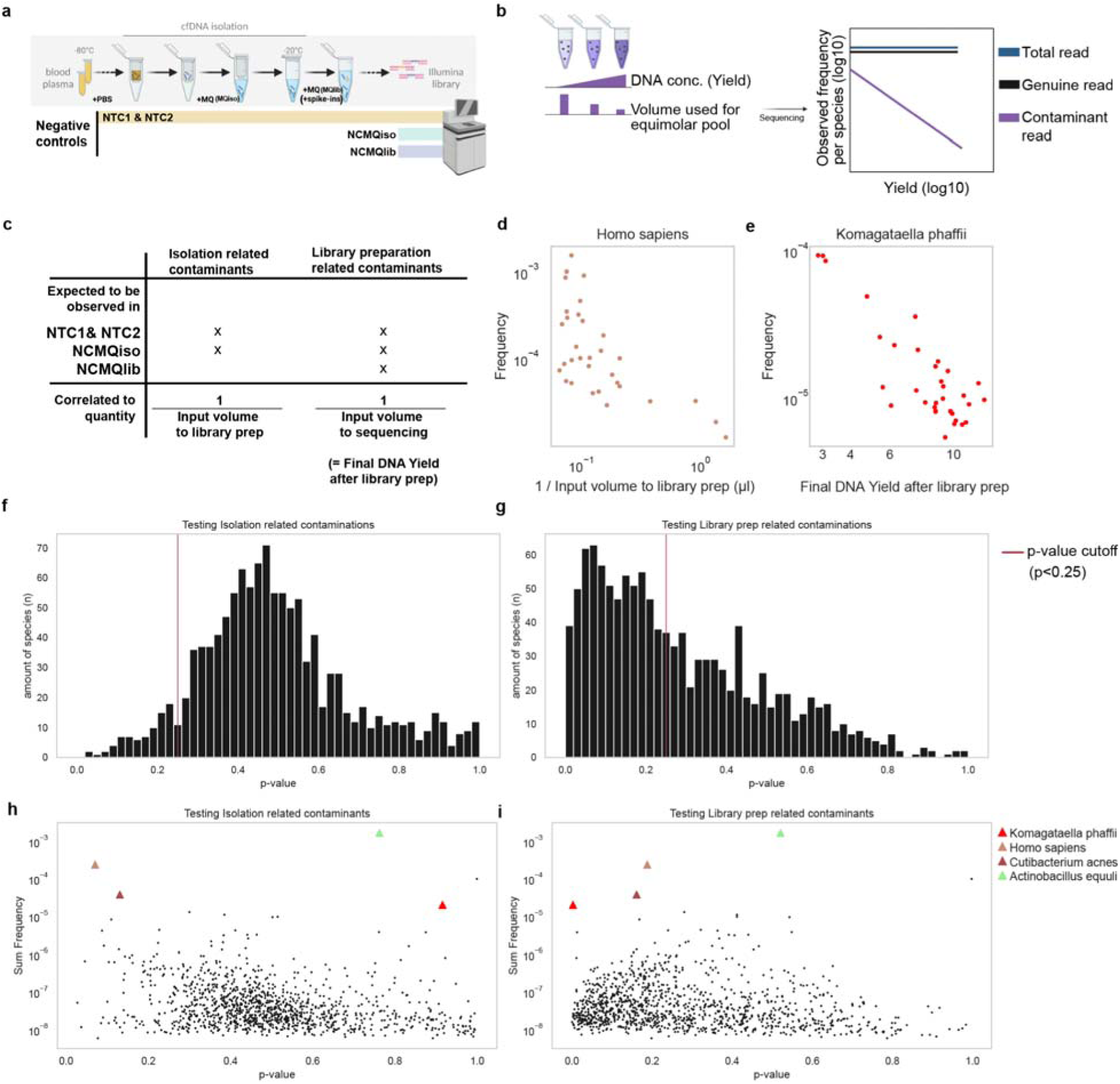
Identification of contaminant taxonomic species. **a.** Schematic of four negative controls used in our study: Non-Template Controls (NTC1, NTC2), which underwent cfDNA DNA isolation and NGS library preparation, and Nuclease-Free water controls (NC1MQiso, NC1MQlib), which underwent only NGS library preparation. **b.** Contaminants in our samples may have been introduced during cfDNA isolation or sequencing library preparation. Contaminant DNA is expected to be present in low, relatively uniform concentrations across laboratory equipment and kits, leading to similar levels across samples. In contrast, cfDNA concentration - and thus the yield from cfDNA isolation or library preparation - can vary significantly between samples. Consequently, the expected frequency of contaminant DNA decreases as the total DNA (yield) in the sample increases (purple), while the frequency of non-contaminant DNA remains consistent (black). Figure adapted from (Davis et al. 2018) **c.** Table presents the expected cfDNA isolation and library preparation contaminants in each negative control sample (top), along with how input volumes serve as an inverse proxy for yield (bottom). **d.** Dotplot showing th relative frequency of *Homo sapiens* reads, an identified cfDNA isolation-related contaminant, versus the input volume used for library preparation (inverse proxy for cfDNA yield) across 32 samples. **e.** Dotplot showing the relative frequency of *Komagataell phaffi* reads, a proven library preparation-related contaminant, versus the input volume used for pooling (inverse proxy for library preparation yield) across 32 samples. **f.** Histogram of p-values derived from the decontam method for cfDNA isolation-related contaminants in all species tested (n = 1167). The p-value detection classification threshold is set at 0.25, indicated by the red vertical line; species falling below this threshold are identified as cfDNA isolation-related contaminants. **g.** Histogram of p-values derived from the decontam method for library preparation-related contaminants in all species tested (n = 1167). The p-value detection classification threshold is set at 0.25, indicated by the red vertical line; species falling below this threshold are identified as library preparation-related contaminants. **h.** Scatter plot displaying the summed frequency across all foals against p-values derived from the decontam method for cfDNA isolation-related contaminants. Contaminants *Homo sapiens* (p < 0.25) and *Cutibacterium acnes* (p < 0.25) are highlighted, indicated by red and brown triangles, respectively. The NGS library preparation-associated contaminant *Komagataella phaffi* (not significant) is also marked with a triangle. Additionally, *Actinobacillus equuli*, a highly abundant microbial species (frequently observed pathogen in previous research) that is not significantly detected as a contaminant, is represented by a green triangle. **i.** Scatter plot displaying the summed frequency across all foals against p-values derived from the decontam method for library preparation-related contaminants, highlighting contaminants. Contaminants *Homo sapiens* (p < 0.25) and *Cutibacterium acnes* (p < 0.25) are highlighted, indicated by red and brown triangles, respectively. The library preparation-associated contaminant *Komagataella phaffi* (p < 0.25) is also marked with a triangle. Additionally, *Actinobacillus equuli*, a highly abundant pathogen that is not significant, is represented by a green triangle.

**Supplementary Figure 4.**
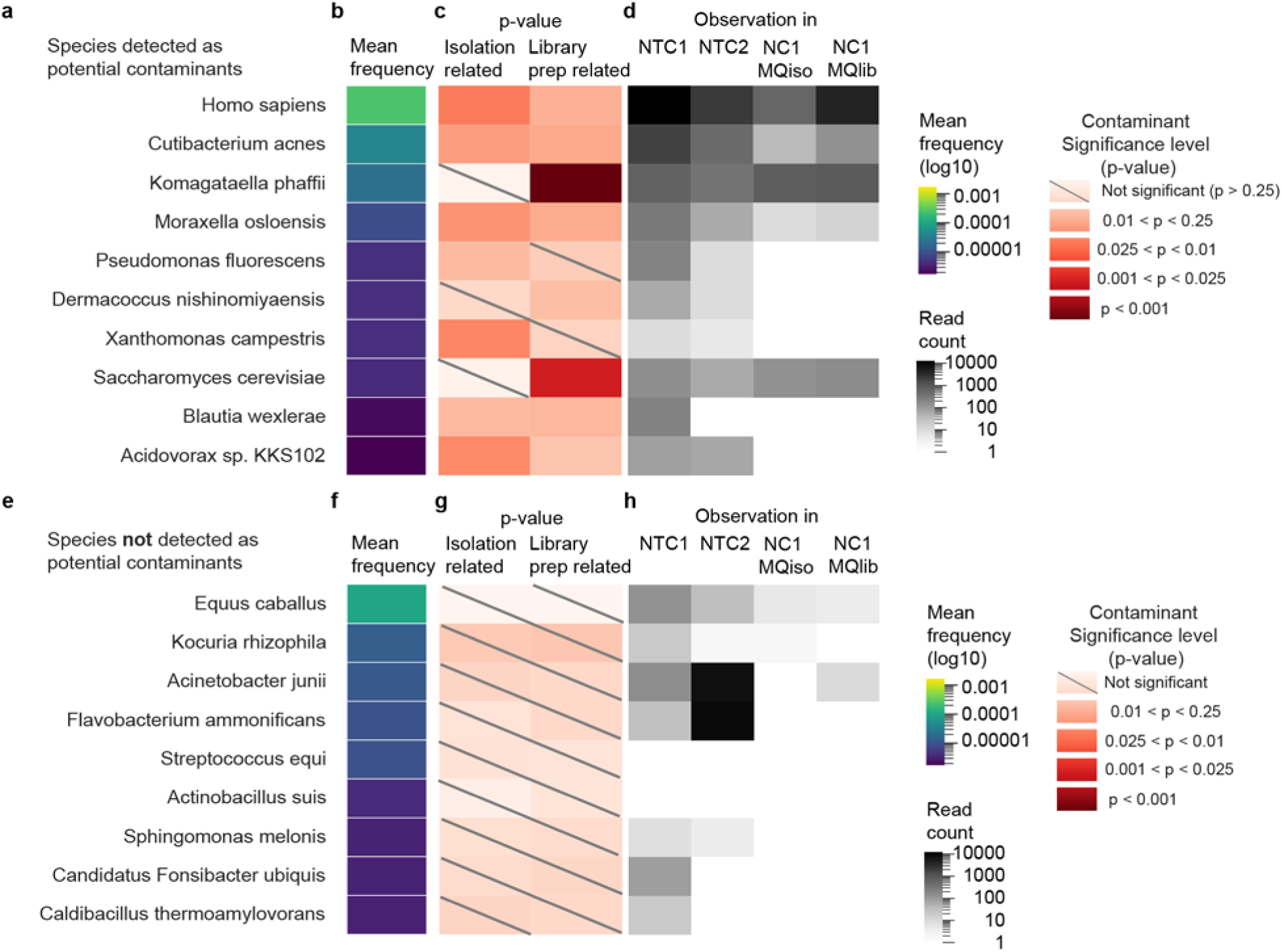
Top species detected as contaminants and not detected as contaminants and their significance and observation in negative controls. **a.** Top 10 species detected as potential contaminants sorted by mean relative abundance. **b.** Heatmap illustrates the mean frequency of potential contaminant species listed in a. The color reflects the log10 mean frequency of each species across all foal samples. **c.** Heatmap displays p-values indicating whether the contaminants are associated with the cfDNA isolation process or the library preparation process. Significance levels range from not significant (light pink, p > 0.25) to highly significant (dark red). **d.** Observations of contaminant species in various negative control conditions: NTC1, NTC2, NC1MQiso, and NC1MQlib. The color intensity reflects the log10 count frequency. **e.** Top 10 species non-contaminant species sorted by mean relative abundance. **f.** Heatmap illustrates the mean frequency of potential non-contaminant species listed in e. The color reflects the log10 mean frequency of each species across all foal samples. **g.** Heatmap of p-values indicating whether the species not detected as contaminants are related to the isolation process or the library preparation process. All were p> 0.25. **h.** Observations of non-contaminant species in various negative control conditions: NTC1, NTC2, NC1MQiso, and NC1MQlib. The color intensity reflects the log10 count frequency.

**Supplementary Figure 5.**
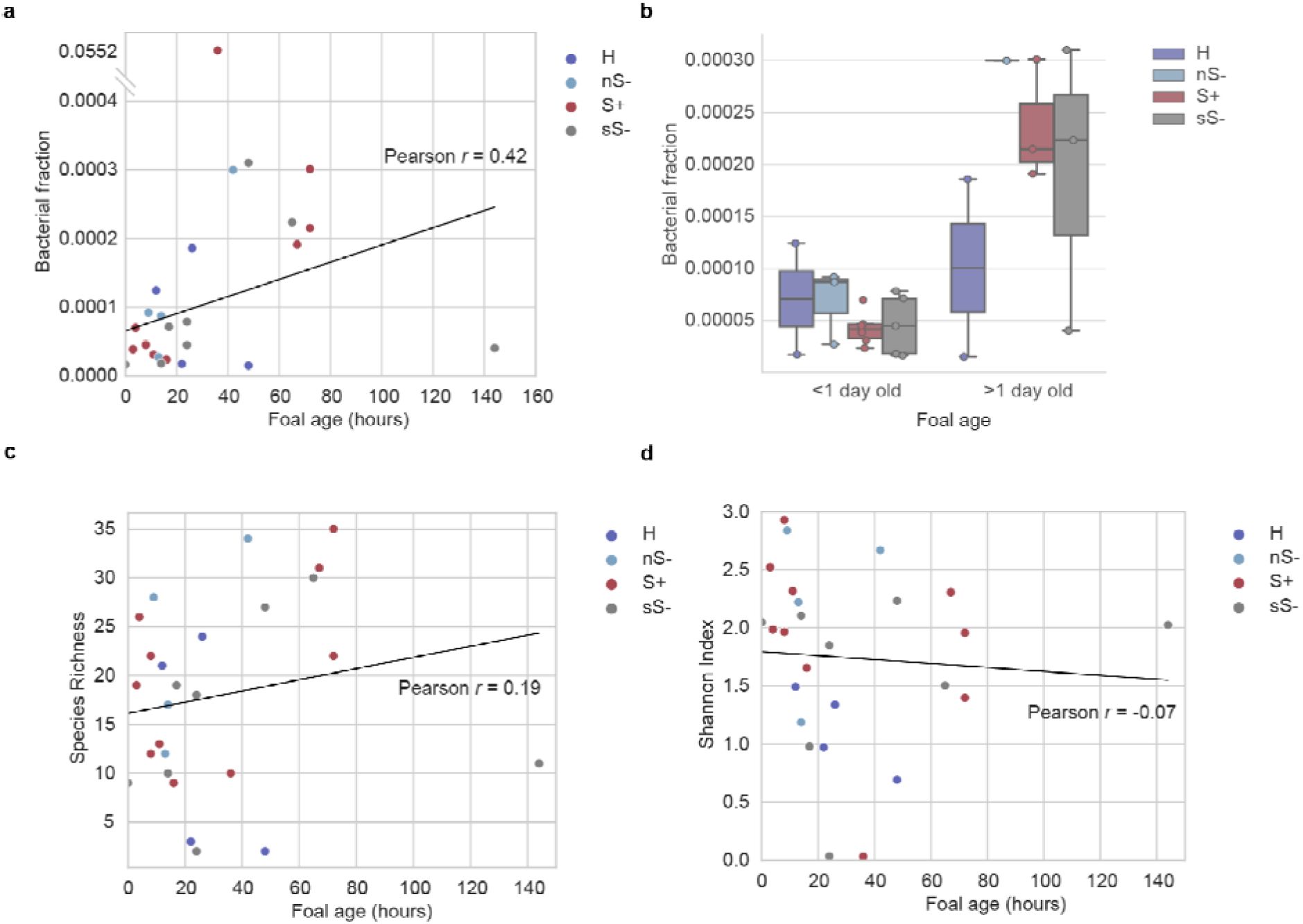
Bacterial fraction, species richness, and Shannon index in foals at various ages, categorized by disease status. **a.** Scatter plot showing the bacterial fraction in relation to foal age at presentation (hours). Data points are color-coded based on disease status. A weak Pearson correlation (*r*=0.42) was calculated excluding outlier data point at bacterial fraction = 0.0552 to avoid strong influence on fitting caused by the outlier. If including this data point, the pearson correlation is *r*=0.03. **b.** Box plot illustrates the bacterial fraction in foals less than one day old compared to those more than one day old. Data points are grouped by health status as shown in a., with whiskers extending to the rest of the distribution within 1.5 times the interquartile range. **c.** Scatter plot depicting species richness in relation to foal age at presentation (hours). Data points are color-coded by health status as in a.. Weak correlation was observed (Pearson correlation *r*=0.19). **d.** Scatter plot showing the Shannon index in relation to foal age at presentation (hours). Data points are color-coded by health status as in a.. Weak negative correlation was observed (Pearson correlation *r*=0.19).

**Supplementary Figure 6.**
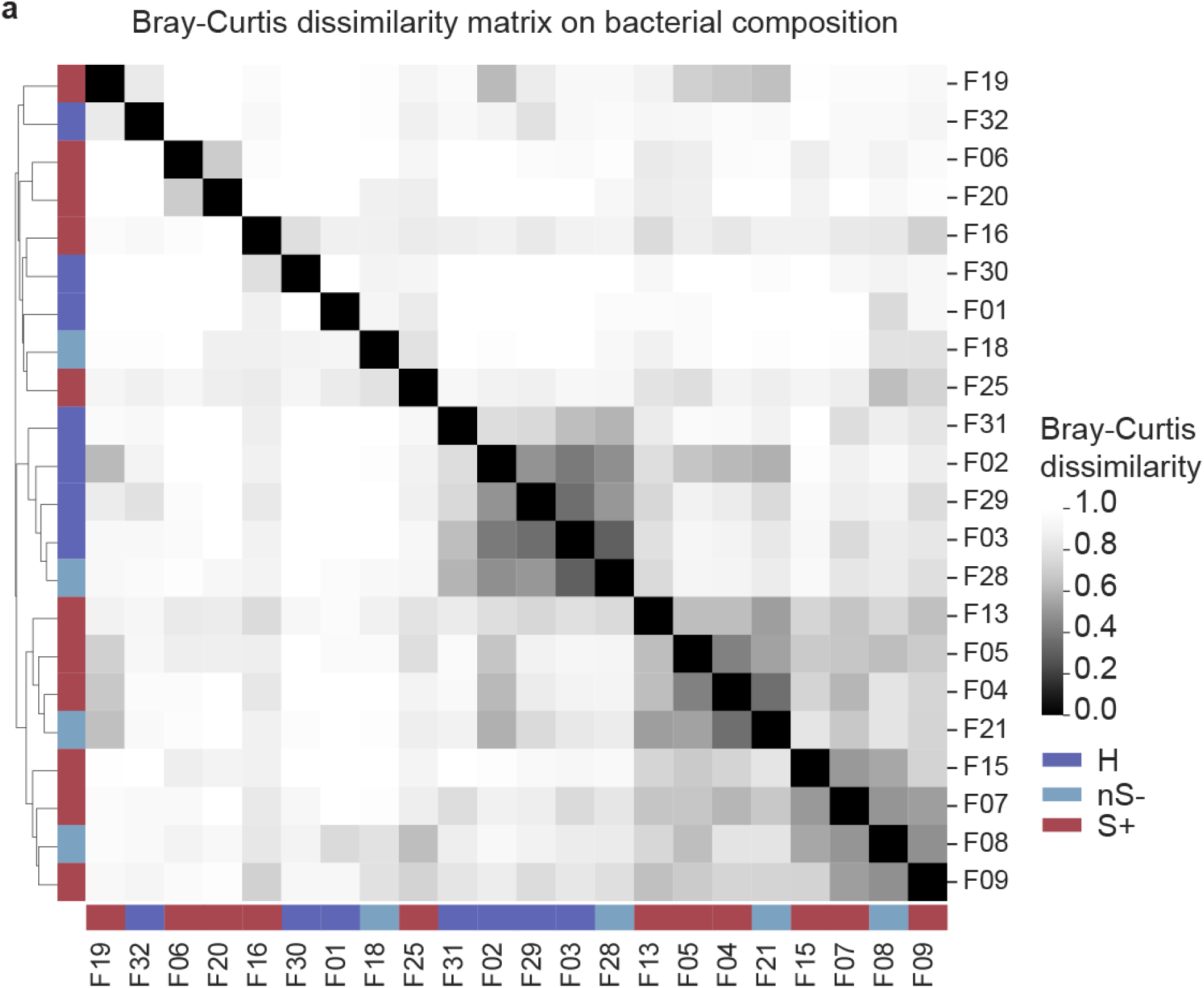
Bray-Curtis dissimilarity matrix showing the distances in log transformed bacterial composition between samples. Distances are calculated for each pair of foal plasma samples, and the samples are clustered based on the minimum linkage of euclidean distances of pairwise Bray-Curtis dissimilarity.

**Supplementary Figure 7.**
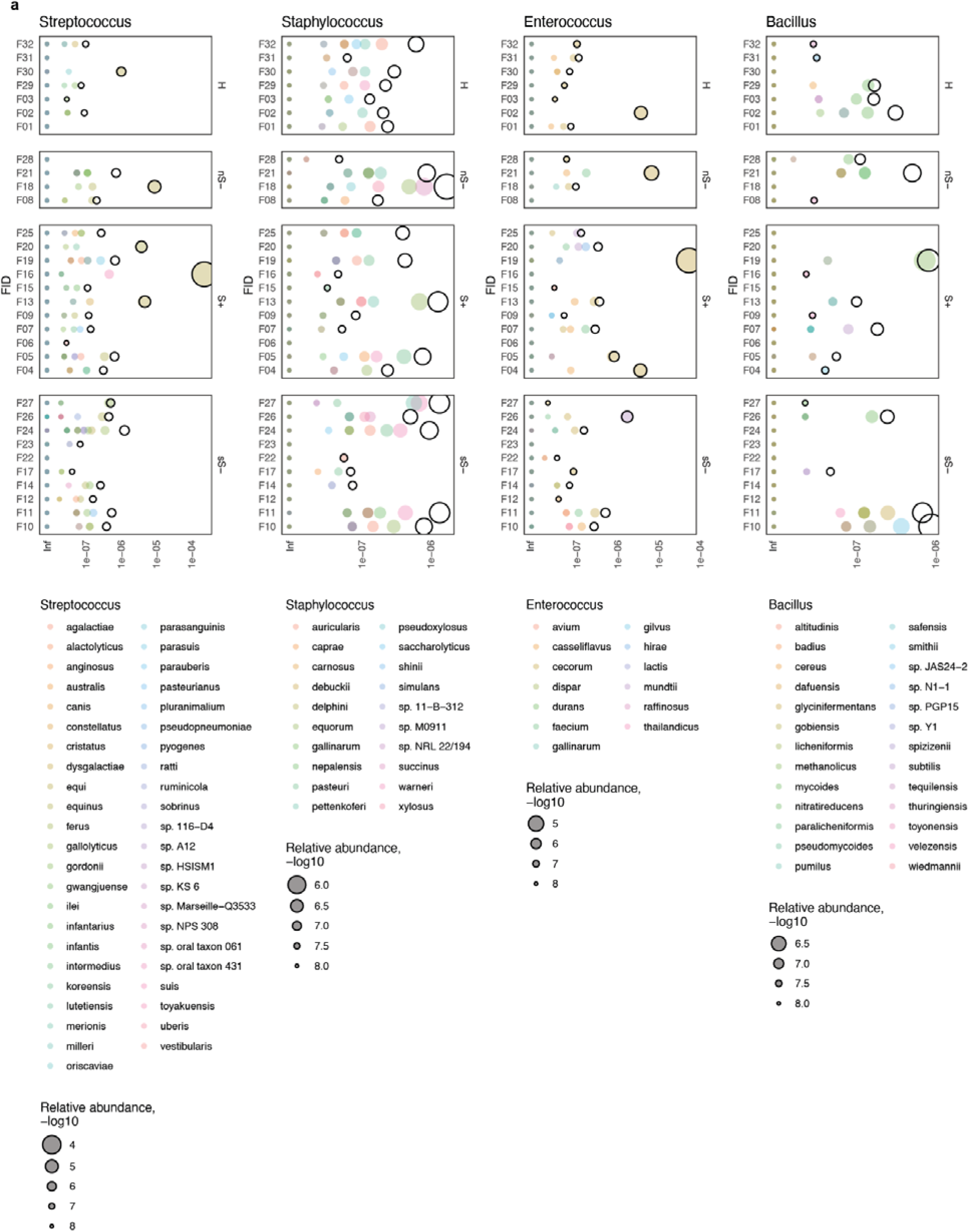
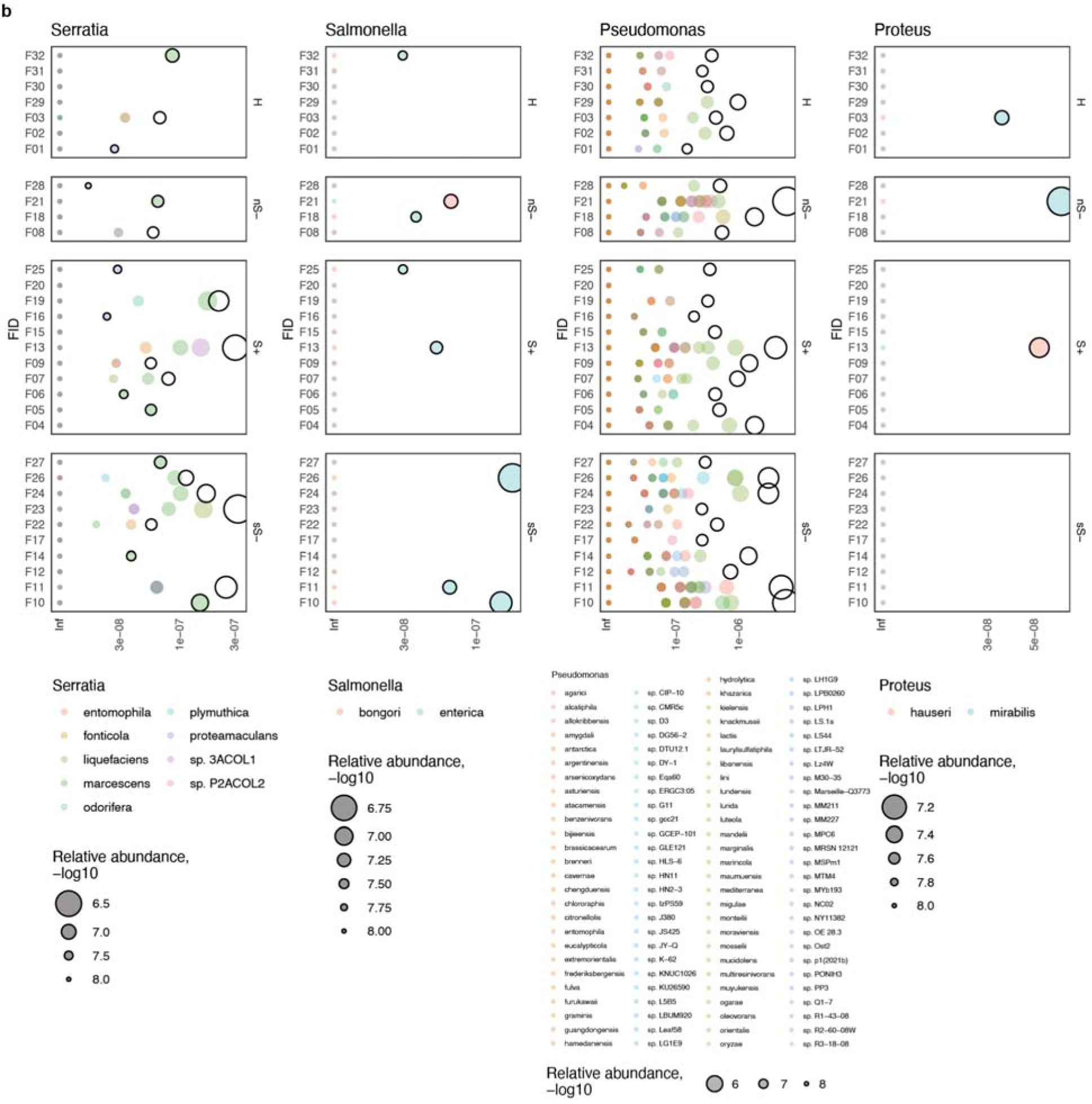
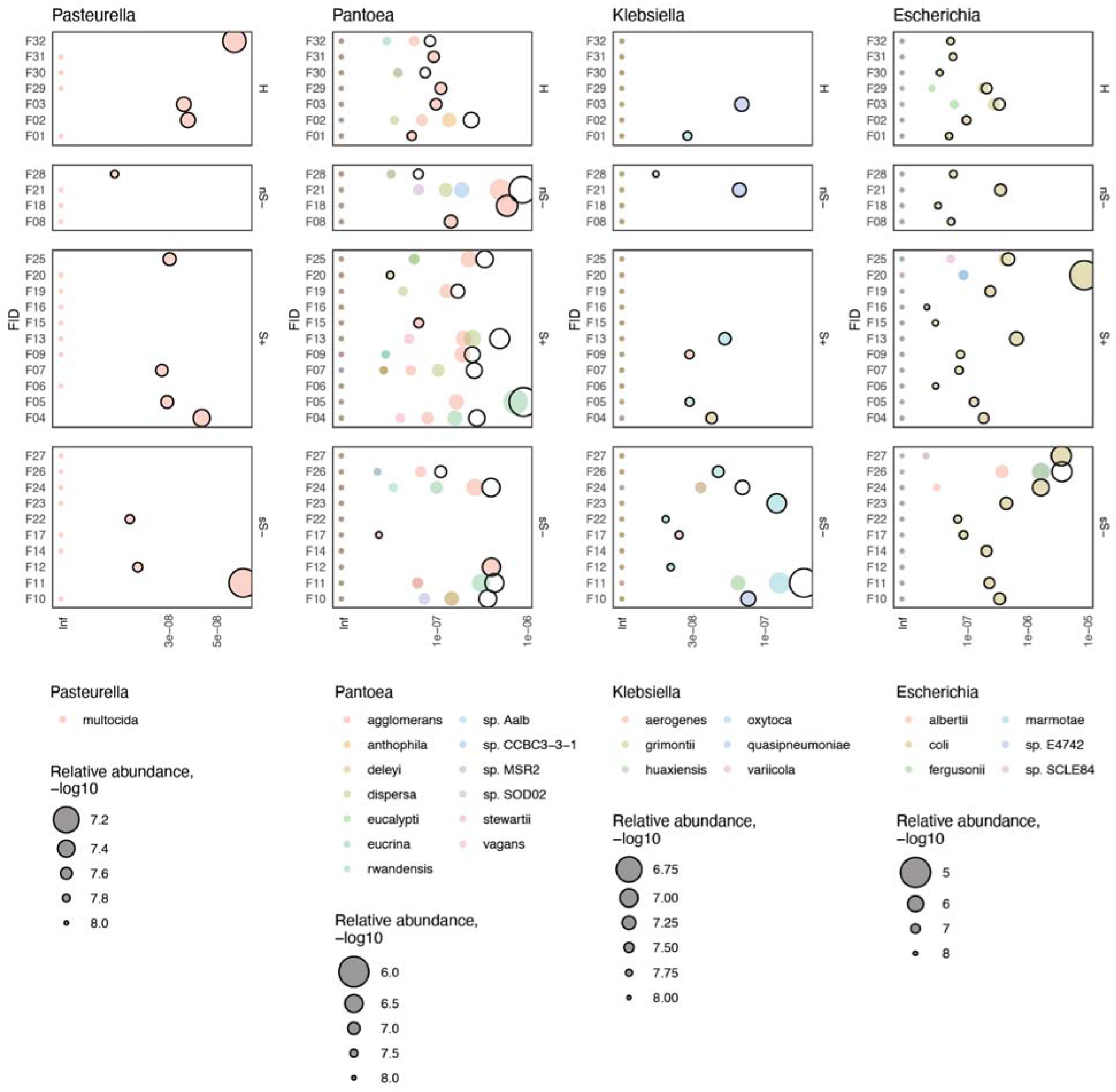
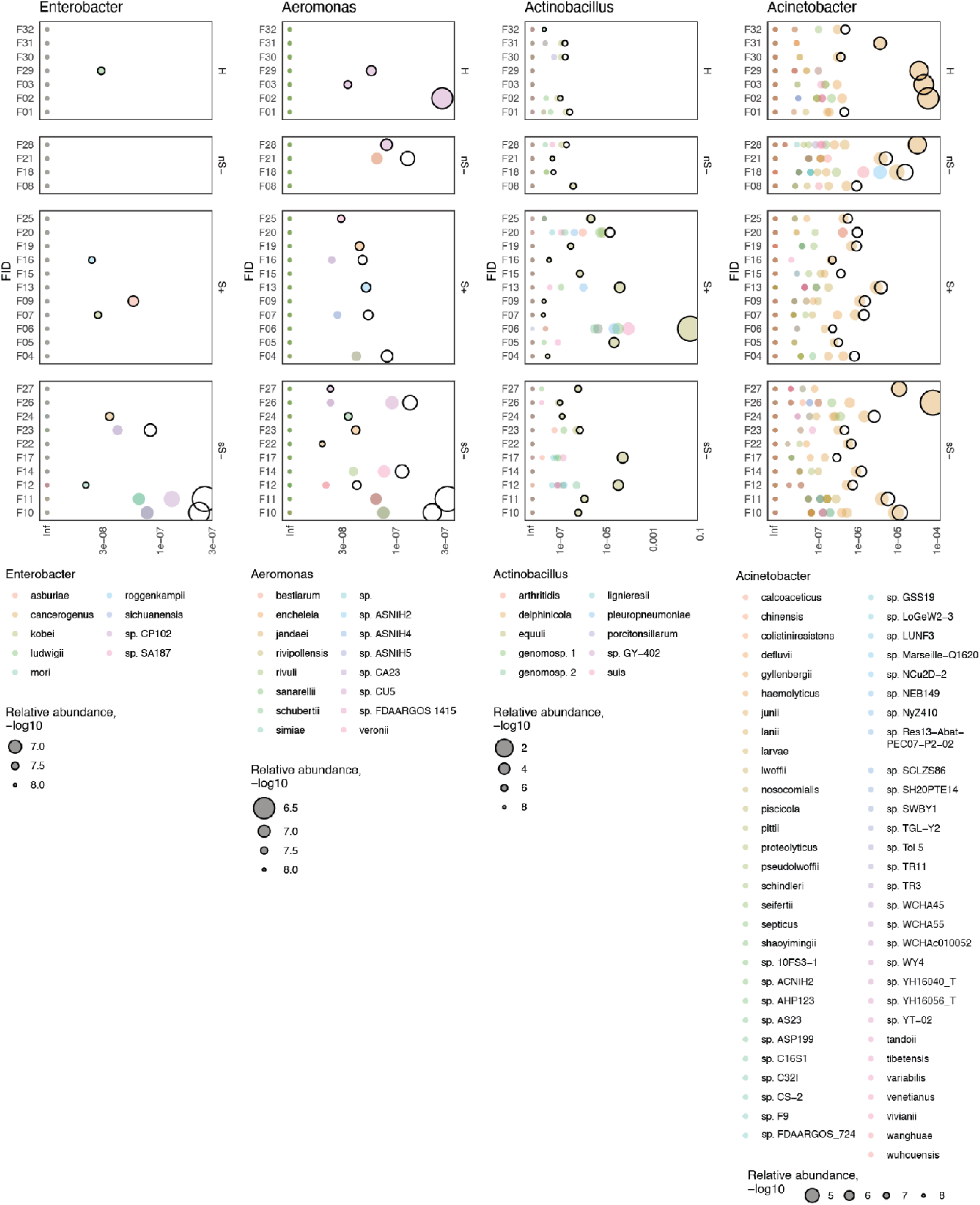
Relative abundance of pathogenic bacterial species and genera. **a.** The dot plot illustrates the relative abundance (x-axis) of species from four selected Gram-negative pathogenic bacterial genera. It also shows the total relative abundance of these genera (sum of species) across samples (represented by black circles). Each dot color represents a different species, while the dot size indicates their relative abundance. For species with no observations, relative abundance is indicated as infinity (Inf). **b.** The dot plot illustrates the relative abundance (x-axis) of species from four selected Gram-negative pathogenic bacterial genera. It also shows the total relative abundance of these genera (sum of species) across samples (represented by black circles). Each dot color represents a different species, while the dot size indicates their relative abundance. For species with no observations, relative abundance is indicated as infinity (Inf).

**Supplementary Figure 8.**
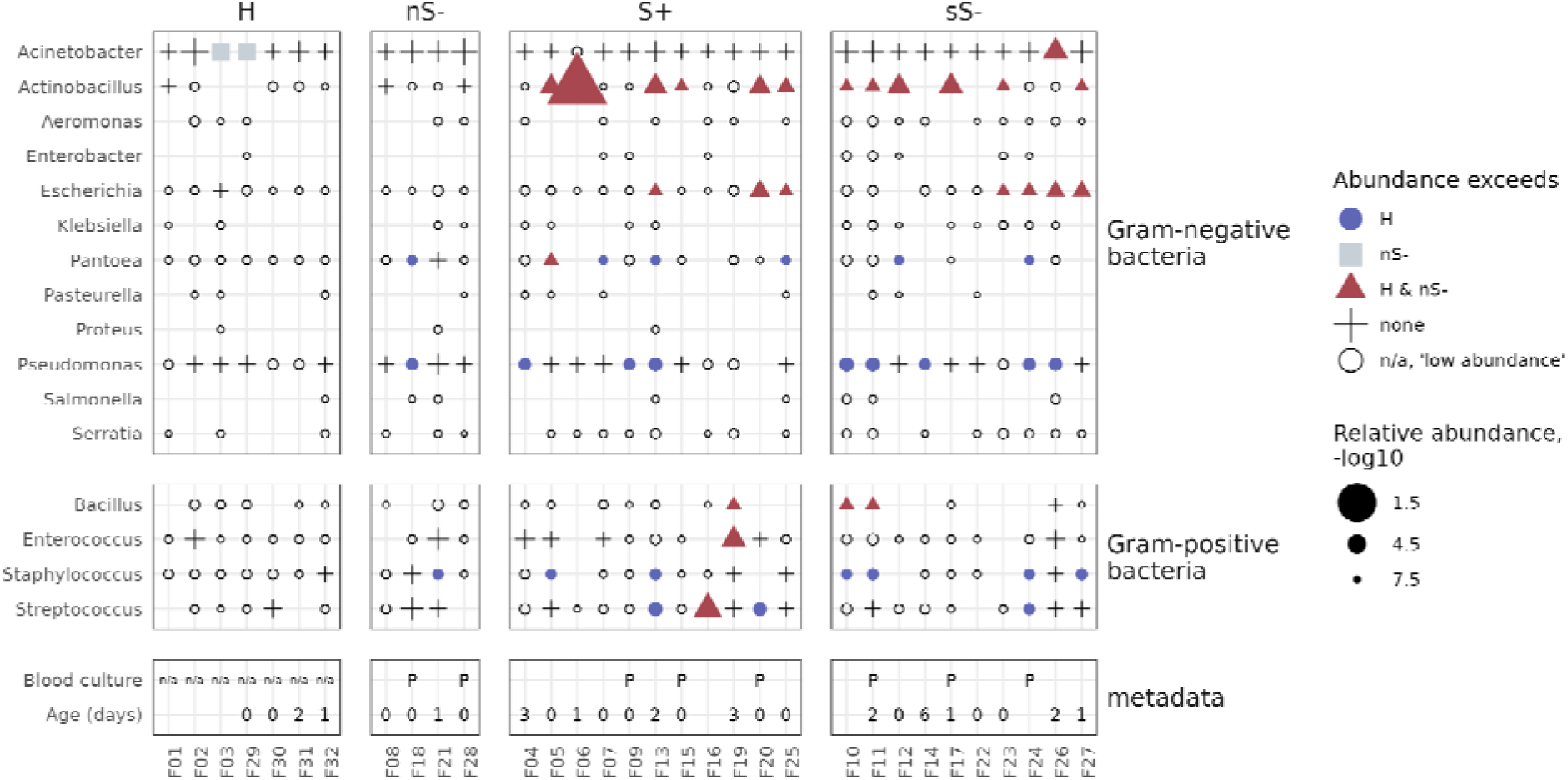
Detection of pathogenic bacteria in sick foals. Dotplot displaying the detection of the 16 most frequently cultured pathogenic genera. Gram-negative species are shown at the top, and Gram-positive species are shown in the middle. Each dot on the plot represents a genus that was detected with at least 1 read. The different symbols and colors convey the relative abundance of the genera in comparison to healthy (H) foals and nS- foals: Blue circles indicate genera with a relative abundance higher than in H foals. Light-gray squares indicate genera with a relative abundance higher than in nS- foals. Red triangles denote genera with a relative abundance higher than in H and nS- foals. White circles represent genera detected at low abundance (fewer than 10 reads) and therefore not compared to either H or nS- foals. Plus signs (‘+’) represent genera that are not elevated compared to either healthy (H) or non-suppurative (nS-) foals. Metadata i displayed at the bottom, including blood culture results (‘P’ for positive, ‘n/a’ for not performed) and age at hospital presentation, if known.

**Supplementary Figure 9.**
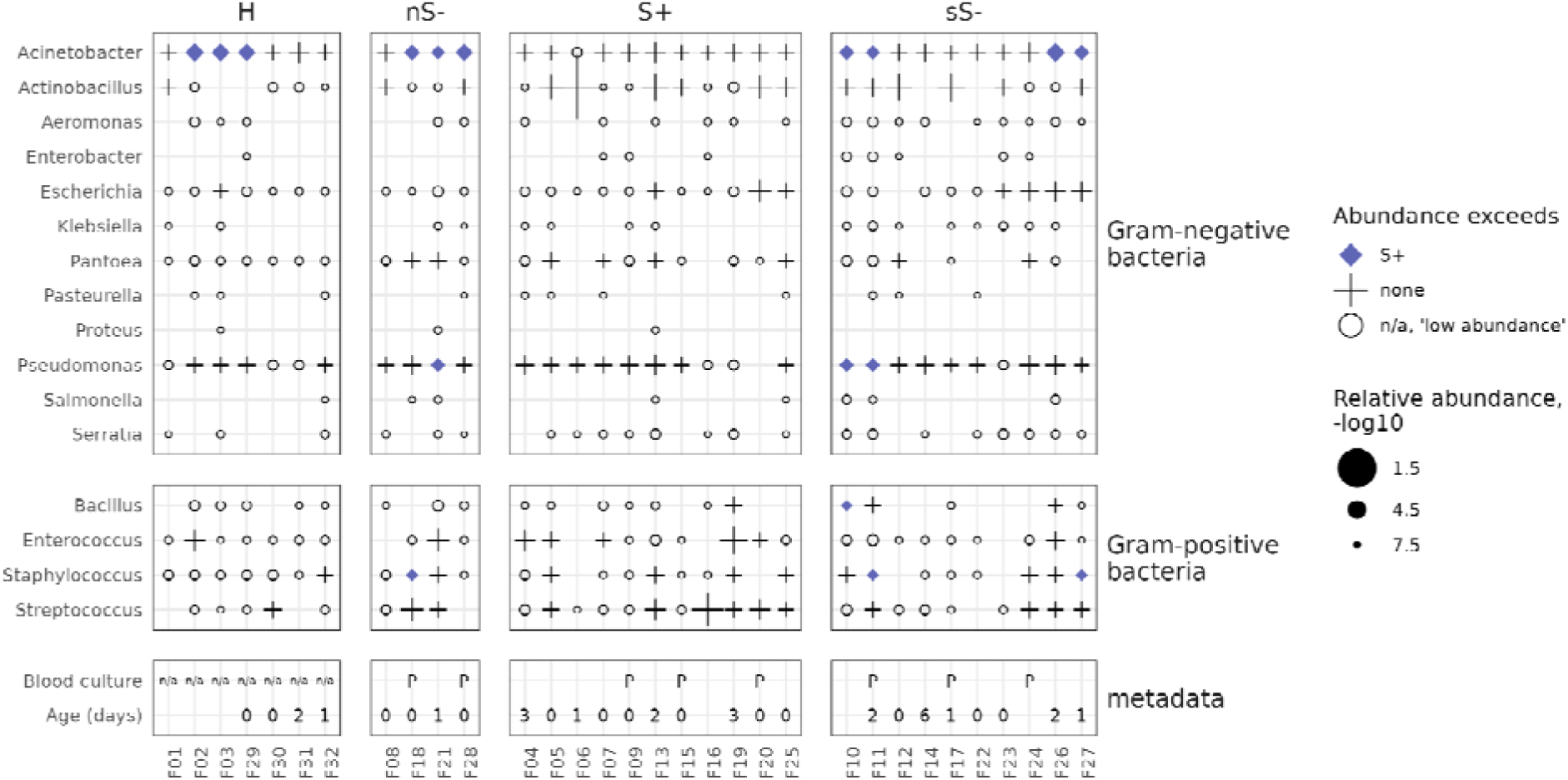
Relative abundance of pathogenic bacterial species and genera compared to S+. Dotplot displaying the detection of the 16 most frequently cultured pathogenic genera. Gram-negative species are shown at the top, and Gram-positive species are shown in the middle. Each dot on the plot represents a genus that was detected with at least 1 read. The different symbols and colors convey the relative abundance of the genera in comparison to healthy (H) foals and nS- foals: Blue diamonds indicate genera with a relative abundance higher than in S+ foals. White circles represent genera detected at low abundance (fewer than 10 reads) and therefore not compared to either S+ foals. Plus signs (‘+’) represent genera that are not elevated compared to S+ foals. Metadata is displayed at the bottom, including blood culture results (‘P’ for positive, ‘n/a’ for not performed) and age at hospital presentation, if known.

**Supplementary Figure 10.**
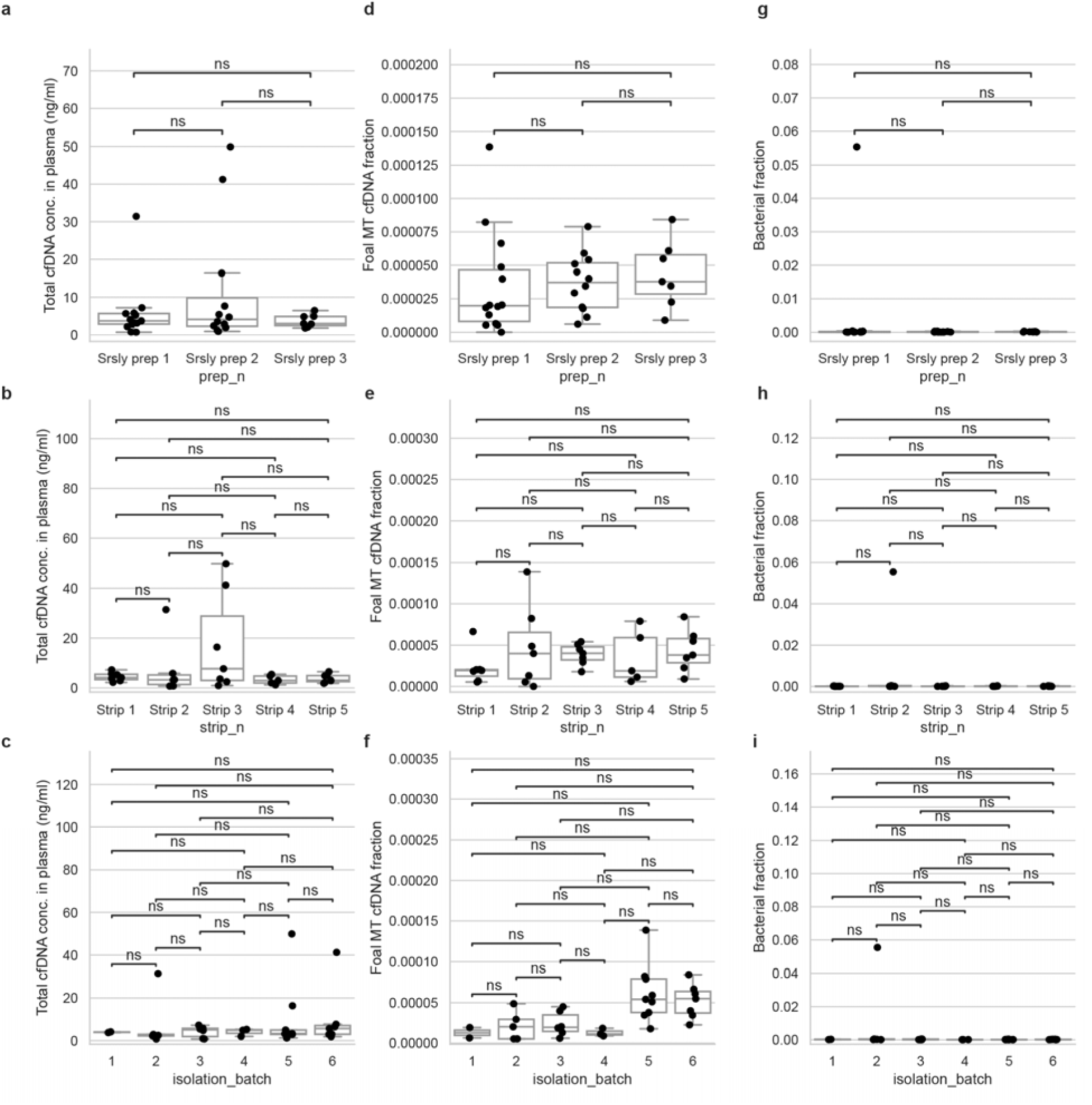
Influence of various batch effects on total cfDNA concentration in plasma, foal mitochondrial cfDNA fraction, and bacterial fractions. Batch effects include different sample preparation methods, strip variations, and isolation batches. Each box plot displays raw data points and represents the 25th percentile (bottom), median (middle), and 75th percentile (top), with whiskers extending to the rest of the distribution within 1.5 times the interquartile range. A Mann-Whitney U Test with Bonferroni correction was performed, revealing no significant differences (ns) among all tests, indicating the robustness and reproducibility of the cfDNA measurement procedures. **a.** Total cfDNA concentration in plasma (ng/mL) for three different library preparation batches, SRSLY prep 1, SRSLY prep 2, and SRSLY prep 3. **b.** Total cfDNA concentration in plasma (ng/mL) across five different strip locations (Strip 1 to Strip 5). **c.** Total cfDNA concentration in plasma (ng/mL) across six isolation batches. **d.** Fraction of MT cfDNA for the three library preparation batches. **e.** Fraction of MT cfDNA across the five strip locations. **f.** Fraction of MT cfDNA across the six isolation batches. **g.** Bacterial fraction of cfDNA for the three library preparation batches. **h.** Bacterial fraction of cfDNA across the five strip locations. i. Bacterial fraction of cfDNA across the six isolation batches.

